# Ligand binding and conformational changes of SUR1 subunit in pancreatic ATP-sensitive potassium channels

**DOI:** 10.1101/283440

**Authors:** Jing-Xiang Wu, Dian Ding, Mengmeng Wang, Yunlu Kang, Xin Zeng, Lei Chen

**Author notes:** **Correspondence:** Lei Chen.

## Abstract

ATP-sensitive potassium channels (K_ATP_) are energy sensors on the plasma membrane. By sensing the intracellular ADP/ATP ratio of β-cells, pancreatic K_ATP_ channels control insulin release and regulate metabolism at the whole body level. They are implicated in many metabolic disorders and diseases and are therefore important drug targets. Here, we present three structures of pancreatic K_ATP_ channels solved by cryo-electron microscopy (cryo-EM), at resolutions ranging from 4.1 to 4.5 Å. These structures depict the binding site of the antidiabetic drug glibenclamide, indicate how Kir6.2 N-terminus participates the coupling between the peripheral SUR1 subunit and the central Kir6.2 channel, reveal the binding mode of activating nucleotides, and suggest the mechanism of how Mg-ADP binding on nucleotide binding domains (NBDs) drives a conformational change of the SUR1 subunit.

## INTRODUCTION

ATP-sensitive potassium channels (K_ATP_) are ion channels that selectively allow potassium ions to permeate the cell. Their channel activities are tightly regulated by endogenous nucleotide metabolites. Specifically, they are inhibited by ATP and activated by Mg-ADP. By sensing the intracellular ADP/ATP ratio, K_ATP_ channels tune the potassium ion efflux across the plasma membrane and adjust the membrane potential. Therefore, K_ATP_ channels convert the cellular metabolic status into electrical signals, a unique output that has broad physiological effects. K_ATP_ channels are widely distributed in many tissues, including the pancreas, brain, heart, and smooth muscle (Hibino et al., 2010), and they play important roles in many physiological processes, such as hormone secretion and vasodilatation (Hibino et al., 2010). Genetic mutation of genes that encode K_ATP_ channel subunits can lead to several metabolic diseases and neuronal diseases (Ashcroft et al., 2017; Hibino et al., 2010). Therefore, K_ATP_ channels are important drug targets. Clinically relevant sulfonylureas drugs inhibit pancreatic K_ATP_ channels and serve as insulin secretagogues for the treatment of type II diabetes (Bryan et al., 2005), while K_ATP_ activators, such as potassium channel openers (KCOs), are used for hypoglycemia and show promise for myoprotection (Flagg et al., 2010; Hibino et al., 2010).

Previous studies have established that the functional K_ATP_ channel is a hetero-octamer composed of four inward-rectifying potassium channel 6 (Kir6) subunits and four sulfonylurea receptor (SUR) regulatory subunits (Clement et al., 1997; Shyng and Nichols, 1997; Ueda et al., 1997). Kir6 subunits are encoded by either KCNJ8 (Kir6.1) or KCNJ11 (Kir6.2). Kir6 subunits harbor sites for inhibitory ATP binding. The activities of Kir6 can be enhanced by PIP_2_, which is a signaling lipid present in the inner leaflets of the plasma membrane. SUR subunits are composed of the N-terminal transmembrane domain 0-loop 0 (TMD0-L0) and ATP-binding cassettes (ABC) transporter-like modules. They are encoded by either ABCC8 (SUR1) or ABCC9 (SUR2). SUR subunits bind stimulatory Mg-ADP, KCOs, and inhibitory sulfonylureas and regulate Kir6 channel activity. K_ATP_ channels composed of different isoform combinations are distributed in different tissues and have distinct pharmacological profiles, which are primarily determined by the identity of associated SUR subunits.

The ABC transporter has two nucleotide binding domains (NBDs). Each NBD harbors two adjacent halves of a catalytic site for ATP hydrolysis. Therefore, one NBD must pair with the other NBD to form two functional ATP hydrolysis sites when they are in proximity (Locher, 2016). SUR proteins belong to subfamily C of the ABC family transporters (ABCC). The other two family members of ABCC are multidrug resistance-associated proteins (MRPs) and the cystic fibrosis transmembrane conductance regulator (CFTR). MRP is a transporter with known substrates, and the transporter activity of MRP depends on the hydrolysis of Mg-ATP, whereas the CFTR is an Mg-ATP-gated chloride channel. The hallmark of the ABCC family is that the conserved glutamate residue on NBD1 Walker B motif of canonical ABC transporters is replaced by an aspartate (D855 in SUR1). Therefore, the catalytic activity of one ATP hydrolysis site (degenerate site or nucleotide binding site 1, NBS1) is lost (Aittoniemi et al., 2009), and only one functional ATP hydrolysis site (consensus site or NBS2) remains. The activities of most ABC transporters depend on the hydrolysis of Mg-ATP. However, Mg-ADP is the primary nucleotide species that activates SUR1 to enhance K_ATP_ channel activity (Aittoniemi et al., 2009).

Because of the physiological importance of pancreatic K_ATP_ channels, they have served as a model to study the biochemical and physiological properties of K_ATP_ channels for decades. K_ATP_ channels in pancreatic islets are composed of Kir6.2 channel subunits and SUR1 regulatory subunits. Recently, several structures of pancreatic K_ATP_ channel are solved by us and other groups, including the GBM bound inhibited state (abbreviated, GBM state, EMD-6689) (Li et al., 2017), the ATP+GBM bound inhibited state (abbreviated, ATP+GBM state, EMD-7073 and EMD-8470) (Martin et al., 2017a; Martin et al., 2017b) and the Mg-ATP and Mg-ADP bound state (abbreviated, Mg-ATP&ADP state, EMD-7338 and EMD-7339) (Lee et al., 2017). These structures reveal the architecture of K_ATP_ channels, the details of how different subunits are assembled together, the binding site of sulfonylureas on SUR1, ATP on Kir6.2, and Mg-ADP/ATP on SUR1. However, the mechanisms of the sulfonylureas inhibition and Mg-ADP activation are still lacking.

In this paper, we present three structures of the pancreatic K_ATP_ channel solved using an SUR1-Kir6.2 fusion construct. The first structure is the K_ATP_ channel in complex with ATPγS and GBM, and it is solved to a resolution of 4.3 Å (abbreviated, ATP+GBM state). The second structure is the K_ATP_ channel in complex with ATPγS alone, which is solved to 4.4 Å resolution (abbreviated, ATP state). The third structure is the K_ATP_ channel in complex with Mg-ADP, VO_4_^3-^, PIP_2_ and NN414 (Carr et al., 2003), a ligand combination that favors the NBD-dimerized state of SUR1, which is solved to 4.3 Å (abbreviated, Mg-ADP state). This new structural information, together with previously solved structures, provides mechanistic insights into how the pancreatic K_ATP_ channel works.

## RESULTS and DISCUSSION

### A K_ATP_ channel fusion construct for cryo-EM structure determination

To overcome the compositional heterogeneity of the K_ATP_ channel when heterologously coexpressing SUR1 with Kir6.2 (Li et al., 2017), we created a SUR1-Kir6.2 fusion construct, in which the intracellular C-terminus of SUR1 is covalently linked to the N-terminus of Kir6.2 by a linker (Fig. S1A). Fusion construct with a 6 amino acids (6AA) linker was previously used to elucidate the association and stoichiometry of K_ATP_ channels (Clement et al., 1997) and was recently used to solve the structure of K_ATP_ channel in complex with Mg-ATP and Mg-ADP (Lee et al., 2017). To avoid potential artifacts on the structure of the K_ATP_ channel because of limited linker length, we used a flexible linker with 39 amino acids (39AA), which should be sufficient for an 130 Å linear distance when fully extended. Similar to the wild-type channel, the fusion construct can be inhibited by ATP and activated by Mg-ADP in an inside-out patch clamp, which is the characteristic property of K_ATP_ channel (Fig. S1B). However, inhibition by GBM is markedly decreased (Fig. S1B). Because it has been reported that the individually expressed SUR1 subunit is sufficient for high affinity GBM binding (Aguilar-Bryan et al., 1995) and our construct has an intact SUR1 subunit, we reasoned that the reduced GBM inhibition is due to insufficient coupling between SUR1 and Kir6.2 of our fusion construct, as discussed later. To be noted, it is reported that 1 µM GBM can fully inhibit the 6AA linker construct in whole cell ^86^Rb^+^ efflux assays (Shyng and Nichols, 1997), this is probably because the GBM can bind SUR1 to counteract the activation effect of intracellular Mg-ADP or Mg-ATP, rather than directly inhibits the 6AA fusion channel. It is also reported that 300 µM tolbutamide can attenuate the currents of 6AA construct in inside-out patch clamp recording (Lee et al., 2017). This is possibly due to the low affinity block of sulfonylurea on Kir6.2 channel (Reimann et al., 1999). Whether the 6AA construct can be directly inhibited by the GBM bound on SUR1 remains to be thoroughly determined.

### Structures of the K_ATP_ channel in the ATP+GBM and ATP states

We aimed to obtain structures of K_ATP_ with and without GBM at higher resolution than our previous GBM state (Li et al., 2017), which can be used to locate the GBM density by direct comparison. Our previous study demonstrated that the K_ATP_ channel displays a large degree of conformational heterogeneity when in complex with the high affinity inhibitor GBM alone (Li et al., 2017). The additional ATP molecule, an endogenous inhibitor of K_ATP_, can further stabilize the Kir6.2 subunit to reduce conformational heterogeneity (Martin et al., 2017b). Therefore, we applied ATPγS, a slowly hydrolyzable and functional ATP analog, to stabilize the structures of the K_ATP_ channel with and without GBM (Schwanstecher et al., 1994a).

The fusion channel protein was purified and then subjected to cryo-EM single particle analysis (Fig. S1C and S1D). The structure of K_ATP_ in complex with ATPγS and GBM was solved at an average resolution of 4.3 Å, slightly higher than the structure in complex with ATPγS alone at a resolution of 4.5 Å (Fig. 1A-1C, Fig. S2 and S3A, and Tables S1 and S2). The overall K_ATP_ channel structure and SUR1 conformation in the ATP+GBM state is almost identical to that in the ATP state based on our density maps (Fig. S3B). Therefore, we focus the discussion on the structure of the ATP+GBM state with a higher resolution, unless stated otherwise.

**Figure 1.**
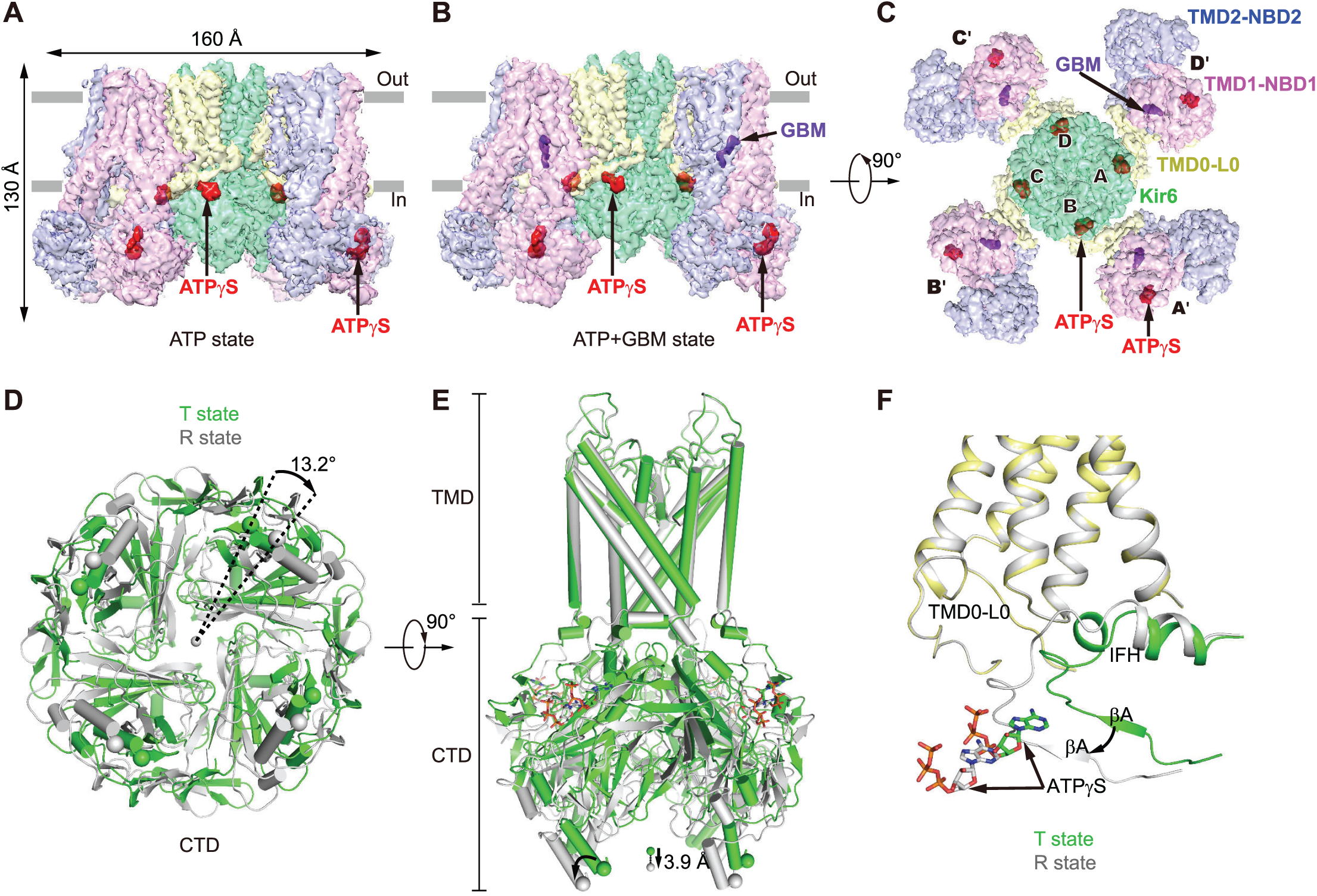
Structures of the pancreatic K_ATP_ channel in complex with ATPγS or ATPγS and glibenclamide (GBM). (A) Side view of the K_ATP_ channel in complex with ATPγS (ATP state). Approximate extent of the phospholipid bilayer is shown as thick gray lines. Kir6.2, SUR1 TMD0-L0 (transmembrane domain 0-loop 0) fragment, TMD1-NBD1 (nucleotide binding domain1), TMD2-NBD2, ATPγS and GBM are shown in green, yellow, magenta, blue, red, and purple, respectively. (B) Side view of the K_ATP_ channel in complex with ATPγS and GBM (ATP+GBM state), colored the same as in (A). (C) Bottom view of the K_ATP_ channel in complex with ATPγS and GBM from the intracellular side. (D) Bottom view of the aligned Kir6.2 structures between “T state” (green) and “R state” (gray) of the ATP+GBM state. Rotation angle between CTDs (cytoplasmic domain) was measured using Cα positions of L356 of Kir6.2, shown as spheres. (E) Side view of the aligned Kir6.2 structures between “T state” (green) and “R state” (gray) of the ATP+GBM state. Vertical movement of CTDs is measured between the centers of mass of L356 Cαs. (F) Positional difference of Kir6.2 βA and IFH (interfacial helix) between “T state” and “R state” structures of the ATP+GBM state.

Focused three-dimensional (3D) classification on the Kir6.2 cytosolic globular domain (CTD) isolated two major classes with distinct conformations (Fig. S2 and Table S1). The CTDs of one class have relative rotation to the other, as observed previously in the structure of the ATP+GBM state (Martin et al., 2017b). Each class was individually refined to reach resolutions of 4.3 Å or 4.6 Å. In contrast, the transmembrane domain (TMD) of the two classes shares the same structure and focused refinement with all particles reached an overall resolution of 4.1 Å (Fig. S4 and Table S1). The maps demonstrate side chain densities for most residues and enable further improvement of our previous model based on medium resolution maps. The overall structures of both classes are similar to the previous symmetric Class 1.1 structure of the GBM state (EMD-6689) (Li et al., 2017) or that of the ATP+GBM state (EMD-8470) (Martin et al., 2017b), with the central Kir6.2 in a closed state and the peripheral SUR1 in an inward-facing inactive state (Fig. 1B and 1C). By comparing the structures of the two classes, we found that the CTDs of Kir6.2 (R32 to L66 and K170 to L356) move in a rigid body fashion relative to the transmembrane domains, resulting in a 3.9-Å downward movement and 13.2°clockwise rotation (Fig. 1D and 1E). In contrast, the structure of transmembrane domain of Kir6.2 and SUR1 is largely unchanged (Fig. 1D and 1E). Therefore, the interface between Kir6.2 CTD and SUR1 TMD0-L0 fragment is altered and the conformations of the linkers between the two rigid bodies also change. Here, we refer to class 1 structure as the “T state” (the tense state) and class 2 as the “R state” (the relaxed state). Specifically, in the “R state” structure, the linker between βA and the interfacial helix (IFH) of Kir6.2 moves away from the TMD0-L0 of SUR1 (Fig. 1F). The differences between the two conformations described herein are not due to differences in ATPγS occupancy on CTDs, because we observed similarly strong ATPγS densities in the maps of both classes (Fig. S4A). Instead, they possibly reflect the endogenous mobility of the CTDs in the Kir6.2-inhibited state, because we observed similar conformational heterogeneity of the K_ATP_ channel in the ATP state and the Mg-ADP state as described later. The co-existence of “T state” and “R state” might be functionally important since the rotation of CTD relates to channel gating in another Kir channel member GIRK2 (Whorton and MacKinnon, 2011, 2013).

### GBM binding site on SUR1

Previous studies demonstrated that mutations of Y230A and W232A on the lasso motif reduce GBM affinity (Vila-Carriles et al., 2007), indicating the lasso motif might be in proximity to or on the binding site of GBM. Therefore, we compared the density maps around lasso motif between the ATP state and the ATP+GBM state, and found that the extra density near the lasso motif in the previous medium resolution map is present in both states. With improved resolution and map quality, we found that this extra density was contributed by both the M5-Lh1 loop and an unknown ligand, probably a lipid head group that constitutively binds K_ATP_, because it is also present in the Mg-ADP state described later (Fig. S4B). Instead, a previously unmodeled density inside the SUR1 central cavity in the ATP+GBM state map has a distinct elongated shape that matches the GBM molecule in an extended conformation (Fig. 2A and Fig. S4C-S4E). In contrast, this density is not present in the ATP state, where GBM was not supplied into the sample (Fig. 2B). This unambiguously demonstrates that the extra density inside SUR1 central cavity in the map of the ATP+GBM state represents the GBM molecule (Fig. 2A and Fig. S4C-S4E). The extended conformation of GBM bound to SUR1 is dramatically different from the compact conformation of GBM bound to human aldo-keto reductase 1C3 subfamily (AKR1Cs) (Zhao et al., 2015) (Fig. S4C). Our results confirmed the GBM binding site reported by another group recently (PDB: 6BAA, EMD-7073) (Martin et al., 2017a). Accompanied by mutagenesis results, they suggested indirect allosteric effects of Y230A or W232A. These mutations on the lasso motif might change the structure of adjacent transmembrane helices that are essential for GBM binding (Martin et al., 2017a). Therefore, mutagenesis results might be affected by allostery, and comparison between the maps of K_ATP_ solved from samples supplied with and without GBM is essential to definitively assign the non-protein density for GBM.

**Figure 2.**
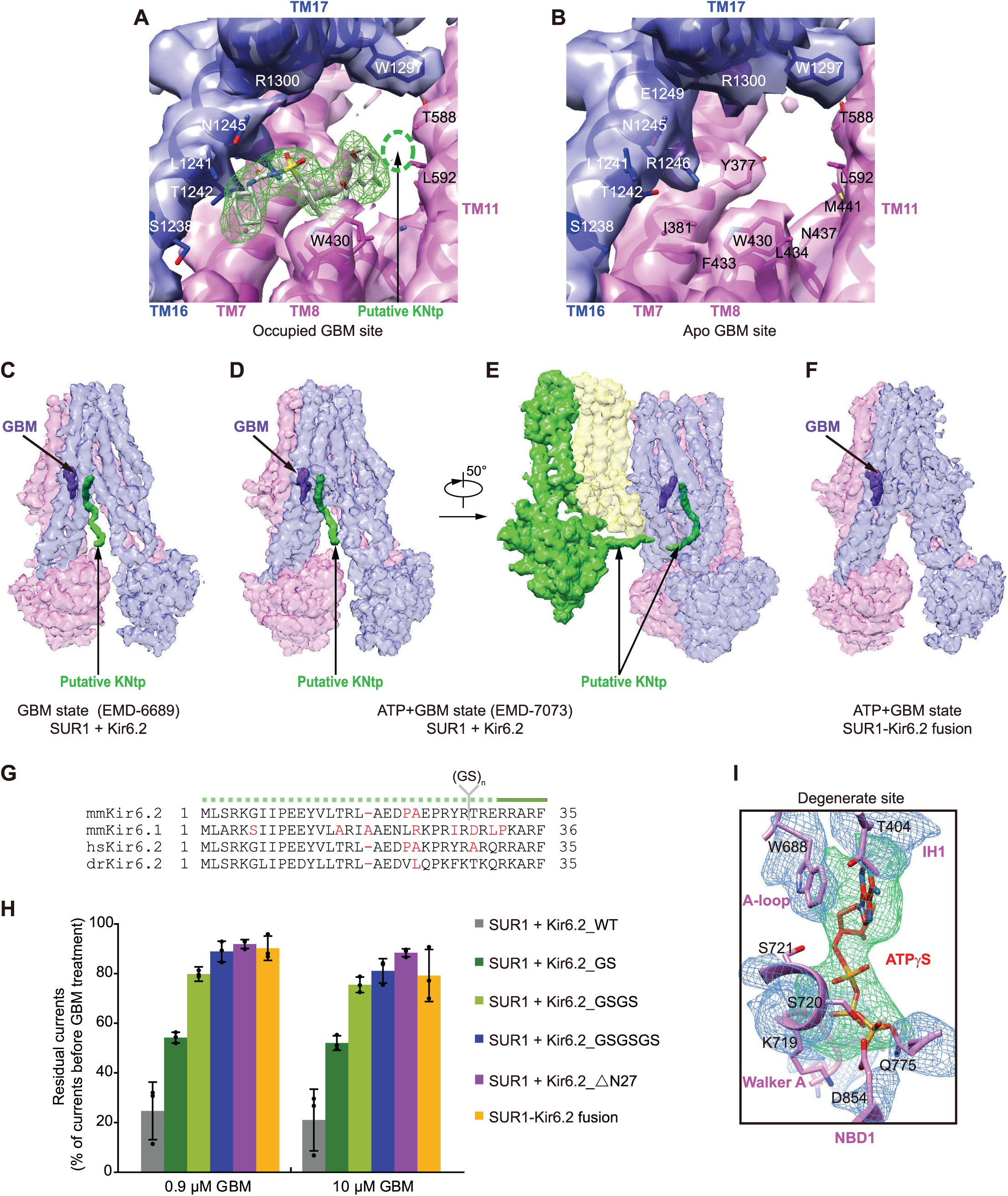
Binding sites of ATPγS and GBM on SUR1. (A) Occupied GBM binding site in the 4.4 Å ATP+GBM state map after focus refinement of SUR1 ABC transporter module. The density of GBM is shown as green mesh with GBM molecule shown in stick representation. The putative KNtp (Kir6.2 N-terminus) binding site is outlined by green dash lines. (B) Apo GBM binding site in the 4.4 Å ATP state map. Maps in (A) and (B) are low-pass filtered to 4.5 Å for comparison. (C-F) EM densities of putative KNtp. Extra continuous density close to GBM molecule is colored in green and GBM density is shown in purple. For better visualization of low resolution features, maps were sharpened with lower B factor than determined by post-processing process. Maps of the GBM state (two half maps for EMD-6689 entry) (C), ATP+GBM state (EMD-7073) (D and E), and ATP+GBM state with SUR1-Kir6.2 fusion construct (F) were sharpened with B factors –200, –50, and +200, and contoured at 0.4, 1.4, and 0.015 levels, respectively. To illustrate the position of KNtp relative to the rest of the Kir6.2 channel, the side view of a single Kir6.2 and SUR1 subunit is shown in (E). (G) Sequence alignment of KNtp of Mus musculus Kir6.2 (mmKir6.2), Mus musculus Kir6.1 (mmKir6.1), Homo sapiens Kir6.2 (hsKir6.2), and Danio rerio Kir6.2 (drKir6.2). Because KNtp is unmodeled due to limited map quality, it is shown as dashed lines above. Nonconserved residues are highlighted in red. Position of the inserted dipeptide (Gly-Ser) is indicated. (H) Comparison of GBM inhibitory effects among Kir6.2 constructs with different numbers of Gly-Ser (GS) dipeptides inserted or N-terminal deletion, when coexpressed with SUR1. The inhibitory effect of GBM on SUR1-Kir6.2 fusion construct is also shown. Currents after GBM treatment were normalized to the currents before GBM treatment. Each bar represents mean ± standard error, n=3. (I) EM densities at the degenerate site of SUR1 in the ATP+GBM state. Densities of ATPγS and surrounding region are shown as green and blue meshes, respectively. Figure is created from map after focused refinement of SUR1 ABC transporter module.

### Location of Kir6.2 N-terminus

The fact that we observed strong GBM density inside SUR1 further reinforced our above assumption that SUR1-Kir6.2 fusion protein can bind GBM but the reduced GBM inhibition is because of insufficient coupling between SUR1 and Kir6.2 due to the covalent linker (Fig. S1B). When docking the improved model of SUR1 and GBM into our previous Class 1.1 map of the GBM state (EMD-6689) (Li et al., 2017), we found an additional positive density in the SUR1 central cavity that originates from the proximity of the GBM molecule and extends out of the SUR1 ABC transporter module in the direction of Kir6.2 CTD (Fig. 2C). Our previous GBM state structure (Li et al., 2017) and the ATP+GBM state structure by another group (Martin et al., 2017b) were solved using K_ATP_ protein generated by the coexpression of SUR1 with Kir6.2 as separated polypeptides; therefore, the Kir6.2 protein had a native N-terminus in these studies. In the map of the ATP+GBM state (EMD-7073), we observed a similar density (Fig. 2D and 2E). Unfortunately, because of limited local map quality, we cannot infer the identity of this density purely from these maps. In contrast, we did not observe any density at the same position in our map of the ATP+GBM state that was solved using the SUR1-Kir6.2 fusion construct, in which the C-terminus of SUR1 is covalently connected to the N-terminus of Kir6.2 by a linker (Fig. 2F) and the linker together with Kir6.2 N-terminus is completely disordered in our cryo-EM map. In our previous models (PDB: 5WUA) (Li et al., 2017), which were built according to the cryo-EM map, only one amino acids of the SUR1 C-terminus were unmodeled, and they are unlikely to contribute to such an elongated unaccounted density. Therefore, this positive density, which is close to the GBM binding site, might represent the N-terminus peptide of Kir6.2 with 30 amino acids omitted in the model. This observation agrees with previous biochemical data showing that Kir6.2 can be labeled by ^125^I-azidoglibenclamide and deletion of KNtp reduced not only ^125^I-azidoglibenclamide labeling (Babenko and Bryan, 2002; Vila-Carriles et al., 2007) but also the affinity for GBM and repaglinide (Kuhner et al., 2012). In addition, GBM enhanced the cross-linking of SUR1 with Kir6.2 with the photo cross-linkable amino acid incorporated in the N-terminus (Devaraneni et al., 2015), indicating that the KNtp should be in proximity to SUR1. This is further supported by the electrophysiological data that high affinity inhibition of tolbutamide, GBM and repaglinide to the K_ATP_ channel were either abolished or markedly decreased when KNtp is deleted (Devaraneni et al., 2015; Kuhner et al., 2012; Reimann et al., 1999). Moreover, the density of the putative KNtp is closed to the 1-chloro-4-methoxybenzene group of GBM (Fig. 2A and Fig. S4E), the so called “B” site, where the azido group in ^125^I-azidoglibenclamide sits. This is in accordance with previous findings that the ^125^I-azidoglibenclamide but not ^125^I-glibenclamide can label Kir6.2 subunit (Aguilar-Bryan et al., 1990; Schwanstecher et al., 1994b). Because the length of KNtp is conserved and limited (Fig. 2G), the special position of the Kir6.2 N-terminus intuitively suggests that it attenuates channel gating by acting as a chain to restrain the rotation of CTD, which is important for Kir family channel gating (Whorton and MacKinnon, 2011, 2013). To further validate this model, we inserted one to three Gly-Ser dipeptides (GS) between amino acid R27 and T28 of Kir6.2, a position before the structured Kir6.2 CTD and not conserved in the primary sequence (Fig. 2G), suggesting this position is not directly involved in SUR1 binding. We hypothesized that the KNtp of these mutants are still able to bind SUR1 but the restriction of the CTD rotation by KNtp might be reduced. Indeed, we found that the inhibitory effect of GBM progressively becomes weaker as the linker length increases (Fig. 2H). The SUR1-Kir6.2 fusion construct demonstrated little GBM inhibition (Fig. 2H and Fig. S1B), a phenotype similar to the K_ATP_ channel assembled with KNtp deleted Kir6.2, suggesting the KNtp in the fusion construct cannot bind SUR1 anymore, because of the covalent linker. Moreover, without sulphonylureas, truncation of KNtp increases channel activity and reduces ATP sensitivity (Babenko et al., 1999; Koster et al., 1999; Reimann et al., 1999) and synthetic KNtp dose-dependently attenuates ATP inhibition (Babenko and Bryan, 2002), suggesting KNtp might remain bound to SUR1 when ATP, but no sulphonylurea, is present, albeit with decreased affinity. When we superposed our SUR1 ABC transporter module model of the GBM+ATP state to that from the another group (PDB: 6BAA) (Martin et al., 2017a), we found that there is a small but noticeable change of overall conformation of SUR1 and the angle between M10 and M16 is 42.7°in our structure whereas 39.3 °in their structure (Fig. S4F). Whether the observed difference of SUR1 conformation is due to KNtp binding discussed above or different data processing procedure still waits further investigation.

### Mg-independent ATP binding site on SUR1 NBD1

During cryo-EM data processing, 3D classification demonstrated continuous conformational heterogeneity of SUR1 in the ATP+GBM state. Specifically, four peripheral SUR1 subunits wobbles around the central Kir6.2 channel in small angles. This greatly limited the resolution of the overall cryo-EM density map and deteriorated local map quality of the peripheral SUR1 ABC transporter module. Further improvement of SUR1 map quality at the ATP+GBM state benefited from focused refinement with local search on symmetry-expanded (Zhou et al., 2015) and signal-subtracted particles of the SUR1 ABC-transporter module (Bai et al., 2015). The map of “the isolated” SUR1 ABC-transporter module reached a resolution of 4.4 Å and demonstrated markedly enhanced quality, especially for NBDs (Fig. S2C). The map also suggests a register shift of residues 1061-1079 on M13 in the structure by another group (PDB: 6BAA) (Martin et al., 2017a). More interestingly, we observed extra density on the degenerate site of SUR1 NBD1. The shape and size of this density matched that of ATPγS (Fig. 2I). Moreover, this density was at the expected location of nucleotides in the nucleotide-bound NBD structure of other ABC transporters (Zhang et al., 2017). Therefore, we modeled this density as an ATPγS molecule. We observed a similar density in the map of the ATP state structure (Fig. 1A). In the sample preparation procedure, we added 2 mM ethylenediaminetetraacetic acid (EDTA) to avoid the activation effect of ATPγS by chelating divalent ions (Proks et al., 2010). Therefore, we observed an Mg-independent ATP binding site in the degenerate site on the NBD1 of SUR1. It is possible that in the ATP+GBM structure reported by another group (Martin et al., 2017a), ATP molecule is also bound on the NBD1, but limited local map quality (EMD-7073) precludes any conclusion.

### Architecture of the K_ATP_ channel in the Mg-ADP bound state

It is previously reported that KNtp is not necessary for channel opening and activation by Mg-ADP (Reimann et al., 1999). The fact that our 39AA fusion construct can be re-activated by Mg-ADP (Fig. S1B) suggests this construct can be used for structural study on the Mg-ADP activation mechanism. To trap SUR1 in an NBD-dimerized conformation, we supplemented the K_ATP_ fusion protein with Mg-ADP, the endogenous activator of the K_ATP_ channel. We also added NN414, a SUR1 specific KCO; VO_4_^3-^, an ion that mimics the post-hydrolytic form of ATP γ-phosphate; and 08:0 PI_(4,5)_P_2_ lipid, a soluble PIP_2_ analogue which is required for K_ATP_ channel activation. The structure of K_ATP_ channel in the ADP bound state was solved to an overall resolution of 4.2 Å (Fig. S5 and Table S3). We observed similar “T state” and “R state” conformational heterogeneity of the Kir6.2 CTD and SUR1 ABC transporter module as in the ATP+GBM state or the ATP state. The map quality of specific regions was dramatically improved by focused classification and refinement (Fig. S5 and Table S3).

The overall architecture of K_ATP_ in the ADP bound state is also similar to the ATP+GBM state or the ATP state (Fig. 3A and 3B). Four Kir6.2 channel subunits are located at the center, and four SUR1 subunits are in the periphery. The SUR1 ABC transporter module connects to the Kir6.2 via SUR1 TMD0-L0 fragment. This is in great contrast to the “quatrefoil form” structure of Mg-ATP and Mg-ADP (Lee et al., 2017) which has a disordered lasso motif and a large rotation of the SUR1 transporter module. The “quatrefoil form” might represent a “decoupled” state, in which the conformational change of SUR1 cannot be transferred to Kir6.2 by either the TMD0-L0 or the N-terminus of Kir6.2. Our overall structure is similar to the coupled “propeller form” (Lee et al., 2017), which has an ordered lasso motif, despite some positional shift of SUR1 subunits (Fig. S6A and S6B). The large difference between the “quatrefoil form” and the “propeller form” might arise from different sample preparation procedures, because detergents were replaced by amphipols in their experiments (Lee et al., 2017). Similarly to the ATP or ATP+GBM state, the CTD of Kir6.2 in “T state” has a clockwise rotation (∼11.8°) relative to that of “R state” (Fig. 3C). The two nucleotide binding sites on SUR1 are occupied by Mg-ADP, which results in an NBD-dimerized conformation (Fig. 3A and 3B). Moreover, the ATP binding sites of Kir6.2 are also occupied by ADP molecules, and the Kir6.2 channel is in the closed conformation, similarly to the ATP+GBM state (Fig. 3A and 3B, and Fig. S6C), which is consistent with the result that ADP can inhibit the K_ATP_ channel albeit at a lower affinity (Hopkins et al., 1992; Schwanstecher et al., 1994a). The ADP binding mode at the inhibitory site of Kir6.2 is similar to that of ATP (Martin et al., 2017a) or ATPγS (Fig. S6C). Because ADP lacks the γ phosphate, there is no direct interaction between ADP and the γ-phosphate-coordinating residue R50 (Martin et al., 2017b). This explains why ADP binds Kir6.2 with a lower affinity and the R50 mutation shifts the IC_50_ curve to a similar range of ADPβS on wild-type channels (Schwanstecher et al., 1994a; Shimomura et al., 2006). We did not observe any density at the PIP_2_ binding site, probably due to the low affinity of the 08:0 PI(4,5)P_2_ to the purified protein and also the negative cooperativity of inhibitory nucleotide binding and PIP_2_ binding on Kir6.2 (Baukrowitz et al., 1998; Hilgemann and Ball, 1996; Shyng and Nichols, 1998). This is in accordance with the fact that we observed a closed state Kir6.2 structure in the Mg-ADP state, because PIP_2_ is necessary for channel opening (Baukrowitz et al., 1998; Shyng and Nichols, 1998).

**Figure 3.**
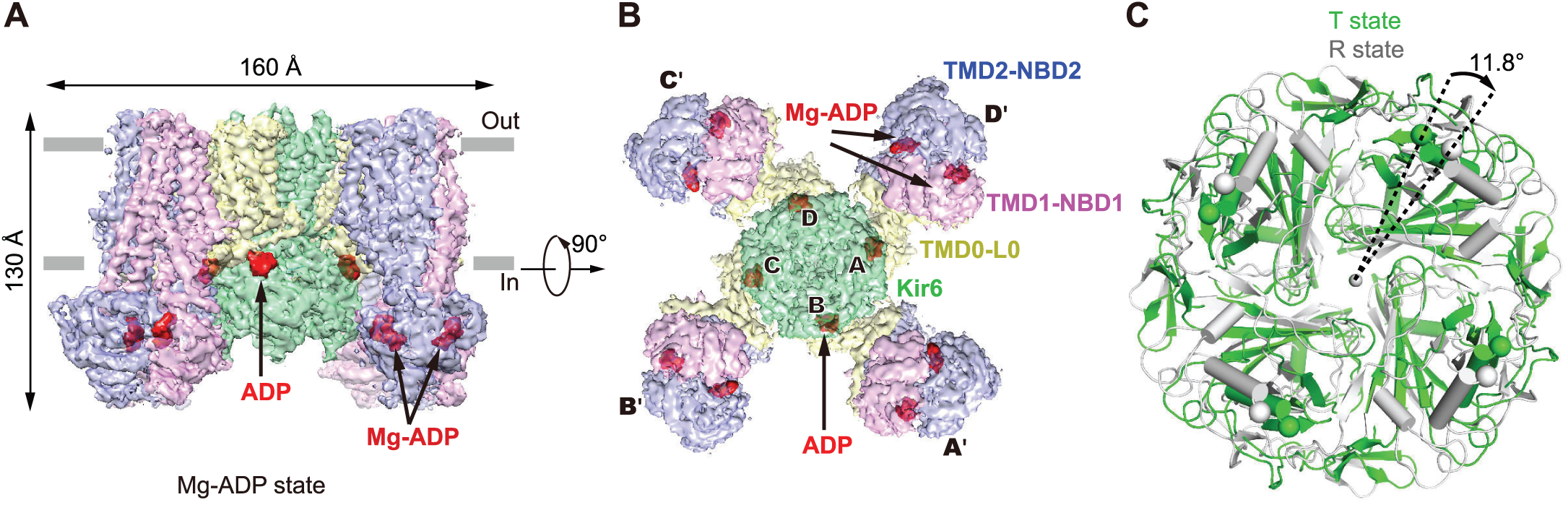
Structure of K_ATP_ in the Mg-ADP state. (A) Side view of the K_ATP_ structure in the Mg-ADP state, colored as in Figure 1A. ADP and Mg-ADP densities are colored in red. (B) Bottom view of the K_ATP_ channel in the Mg-ADP state. (C) Bottom view of the aligned Kir6.2 structures between “T state” (green) and “R state” (gray) of the Mg-ADP state. Rotation angle between CTDs is measured similarly as in Figure 1E.

### Asymmetric NBD dimer of SUR1 in the Mg-ADP bound state

In our focused refined map of SUR1 in the Mg-ADP state, we observed the densities of Mg-ADP molecules in both the degenerate and consensus sites (Fig. 4A-4C). We did not observe densities of VO_4_^3-^, correlating with the fact that VO_4_^3-^ do not markedly affect SUR1 activity (Fig. S1B) (Proks et al., 1999; Shyng et al., 1997; Ueda et al., 1997). The Mg-ADP molecule in the degenerate site interacts with residues from both NBDs to induce a full closure of the degenerate site (Fig. 4C and 4D). In contrast, the Mg-ADP molecule in the consensus site primarily interacts with NBD2, and the consensus site is open (Fig. 4C and 4E). We use the distance between the Cα atoms of the conserved glycine on the ABC Walker A motif and the serine residue (a cysteine on SUR1 degenerate site) on the signature motif as an indicator of NBS closure. The distance between Cα of G1485 and C717 in the degenerate site is 11.5 Å, while the distance between Cα of G833 and S1383 in the consensus site is 13.0 Å (Fig. 4F). These distances are the opposites of the NBD-dimerized state structure of CFTR with Mg-ATP bound (PDB: 5W81) (Zhang et al., 2017). For CFTR, the degenerate site is partially open, and the consensus site is fully closed. The distance between G1350 and S461 in the degenerate site is 12.7 Å, while the distance between G550 and S1249 in the consensus site is 10.6 Å (Fig. 4G). The Mg-ADP state structure has the same asymmetric NBD-dimerized conformation as the Mg-ATP&ADP state structure reported by another group (PDB: 6C3P EMD-7339) (Lee et al., 2017). In particular, Mg-ADP molecule induces similar full closure of degenerate site compared to the Mg-ATP molecules (Fig. S6D).

**Figure 4.**
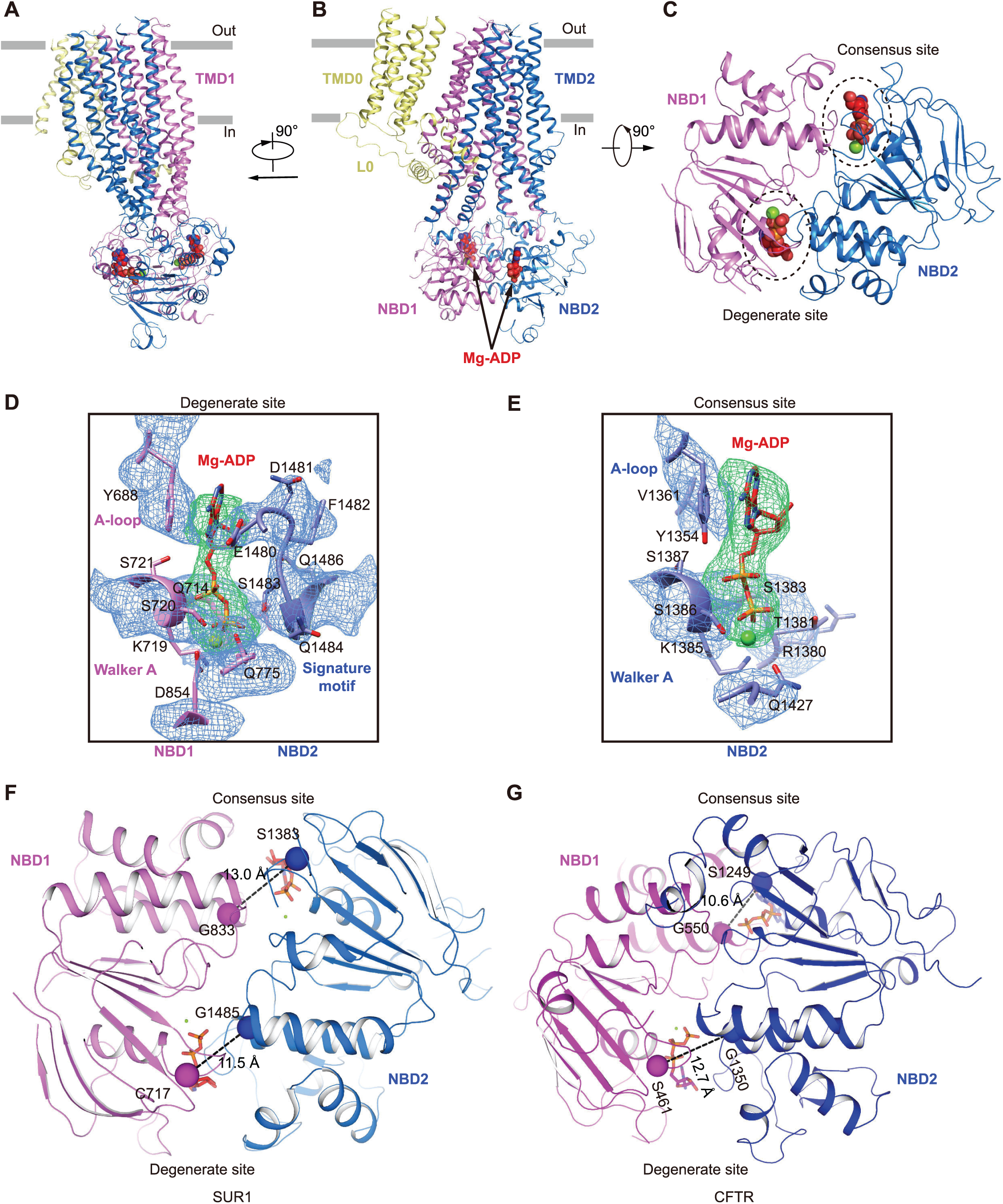
Mg-ADP on SUR1. (A) Side view of a single SUR1 subunit in cartoon representation. Color scheme is the same as in Figure 1A. Mg2+ and ADP are shown as green and red spheres, respectively. (B) Side view of a single SUR1 subunit with a 90° rotation relative to panel (A). (C) Bottom view of a single SUR1 subunit with a 90° rotation relative to (B). For simplicity, only NBD1 and NBD2 are shown. Two Mg-ADP molecules are circled. (D-E) EM densities at the degenerate site (D) and consensus site (E), colored as in Figure 2H. Mg2+ is shown as a green sphere. (F-G) Differences of NBD closures in SUR1 (F) and CFTR (PDB: 5W81) (G). Cα distances between the glycine in the Walker A motif and serine (cysteine instead at the degenerate site of SUR1) in the ABC signature motif are shown as dashed lines.

### Conformational changes of SUR1 ABC transporter module upon Mg-ADP binding

By aligning the TMD0 domain of SUR1 between the ADP and ATP+GBM states, we found a large conformational change of the SUR1 ABC transporter module (Fig. 5A). The full closure of the degenerated site induced by Mg-ADP binding leads to the asymmetric dimerization of two NBDs (Fig. 5B). These conformational changes are further conveyed to the transmembrane domain via coupling helices that interact with NBDs to drive two halves of the transmembrane domain of SUR1 ABC transporter module to move closer (Fig. 5C). Specifically, M14 and M17 move toward M8 and M11 (Fig. 5C). As a result, SUR1 central cavity shrinks, and the GBM and KNtp sites are disrupted. We added NN414, a KCO, to our sample (Fig. S7A). We observed two positive densities in SUR1. One is close to the top of SUR1, surrounded by M10-11 from TMD1, and M12 and M17 from TMD2 (Fig. S7B). The other is at the central cavity surrounded by M8 from TMD1, and M15–17 from TMD2 (Fig. S7C). Both of the densities are absence in the Mg-ATP&ADP state map, where NN414 was not added into the sample (EMD-7338, PDB: 6C3O) (Fig. S7B and S7C). By comparing the structures of transmembrane domain with (Mg-ADP state) and without NN414 (Mg-ATP&ADP state), the ligand corresponding to density 1 introduces local structural changes of M9-11 (Fig. S7D). But whether these densities represent NN414 needs further studies. Notably, we did not observe any density that might account for KNtp. Both our Mg-ADP state structure reported here and the Mg-ATP&ADP state structure reported by another group (Lee et al., 2017) are captured using the fusion construct, where the authentic KNtp is not present. It is possible that the KNtp might adopt an unknown conformation that is different from the one observed in the ATP+GBM state but compatible with our current Mg-ADP state structure.

**Figure 5.**
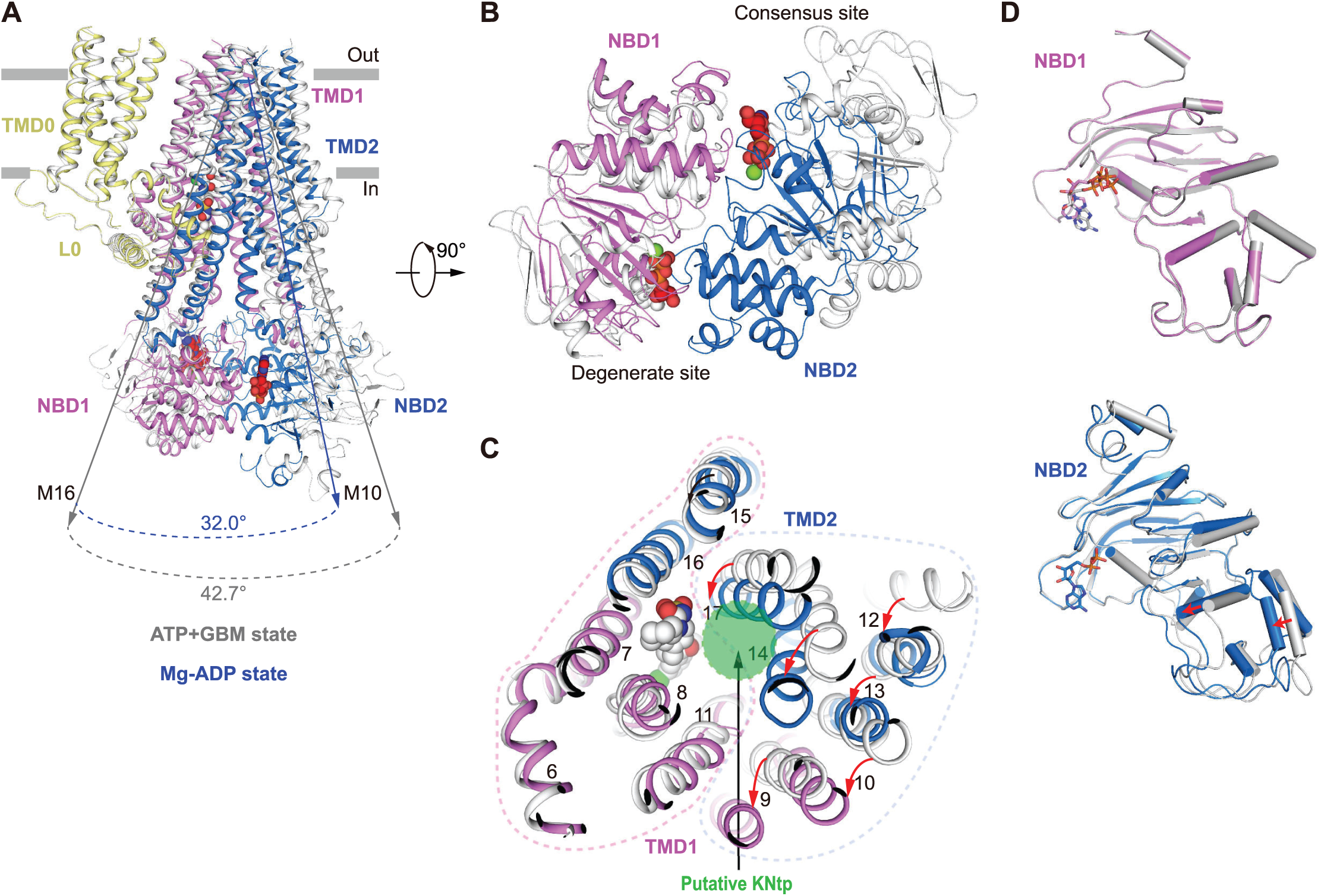
Conformational changes of SUR1. (A) Superposition of the Mg-ADP bound structure (colored) and the ATP+GBM bound structure (gray) by aligning the TMD0 domain of SUR1. Angles between helices M10 and M16 in the two structures are labeled. Mg-ADP, ATPγS, and GBM molecules are all shown as spheres. (B) Bottom view of NBDs of the K_ATP_ channel with a 90° rotation relative to (A). (C) Bottom view of TMDs of the K_ATP_ channel with a 90° rotation relative to (A). Positional changes of transmembrane helices from the first half of TMDs (M9-10, 12-14, and 17) are indicated by red arrows. The dashed lines specify the two halves of TMDs. The putative KNtp binding site is shown as a green circle, showing the displacement of GBM-binding sites upon nucleotide binding. (D) Structural comparisons of individual NBD1 (top) or NBD2 (bottom) between the ATP+GBM state (gray) and the Mg-ADP state (colored). Conformational changes of the α helices-rich subdomain in NBD2 are indicated by red arrows. NBD structures are aligned according to β sheet-rich subdomain.

### Nucleotide binding induced NBD dimerization of SUR1

In the ATP and ATP+GBM state structures, SUR1 is in an inward-facing conformation. We observed ATP density only in the NBD1 degenerate site but not in the NBD2 consensus site, which suggests that the degenerate site has a higher affinity for Mg-free ATP, and ATP alone is not able to drive the dimerization of SUR1 NBDs. This correlates with previous studies that NBD1 of SUR1 can bind 8-azido-ATP with high affinity in an Mg-independent manner (Matsuo et al., 1999a; Matsuo et al., 1999b; Ueda et al., 1997). In addition, the subsequent binding of ATP to NBD1 can be strongly antagonized by prebound Mg-ADP (Ueda et al., 1997), indicating that the degenerate site of NBD1 can be occupied by Mg-ADP as well. Indeed, in our Mg-ADP state structure, we observed Mg-ADP molecules in both the NBD1 degenerate site and the NBD2 consensus site.

The asymmetric NBD dimer conformation observed in the Mg-ADP state structure suggests that the Mg-ADP molecule in the closed degenerate site directly mediates the formation of the NBD dimer interface, because this Mg-ADP molecule physically interacts with both NBDs. In contrast, the Mg-ADP molecule in the open consensus site interacts only with NBD2 but not with NBD1, indicating that it does not contribute directly to NBD dimer formation. By comparing the structures of individual NBD monomer in different states, we found a noticeable conformational change in NBD2 but not in NBD1 (Fig. 5D). Induced conformational changes of the NBD by nucleotide binding were previously observed in the crystal structures of isolated NBD of ABC transporter MJ1267 (Karpowich et al., 2001) and in the cryo-EM structure of CFTR (Zhang and Chen, 2016; Zhang et al., 2017) and also were sampled in MD-simulations (Jones and George, 2017). The conformational changes of NBD2 observed here might partially be due to the Mg-ADP binding and full closure of the degenerate site, but we suggest that the Mg-ADP binding on the NBD2 consensus site plays a more prominent role. This is supported by the previous radioactive 8-azido-ATP photo cross-linking results. First, the NBD2 consensus site mutation K1385M greatly reduces the ability of prebound Mg-ADP to prevent further ATP binding on NBD1, when Mg-ADP and ATP were added to SUR1 sequentially (Ueda et al., 1997; Ueda et al., 1999), suggesting that Mg-ADP acts on the NBD2 consensus site to allosterically increase the affinity of Mg-ADP on the NBD1 degenerate site. Second, Mg-ADP or Mg-ATP binding on the NBD2 consensus site can stabilize prebound ATP on the NBD1 degenerate site (Ueda et al., 1999), indicating that nucleotide binding on the NBD2 consensus site can send a signal to the degenerate site to increase its affinity for nucleotides, probably via the conformational change of NBD2, as we observed here.

### Model for SUR1 conformational change

On the basis of experiments and structures of K_ATP_ channels in complex with different ligand combinations solved with different constructs, we propose the following hypothetic model for the conformational change of SUR1 subunit (Fig. 6A-6H). When GBM is bound, GBM and the Kir6.2 N-terminus cooperatively bind inside SUR1 to inhibit channel gating (Fig. 6A and 6E). Without GBM, the Kir6.2 N-terminus can still bind to the central cavity of SUR1 to inhibit channel gating but with decreased affinity. In the presence of ATP, ATP can occupy the NBD1 degenerate site, but the NBDs are still separated (Fig. 6B and 6F). When the ADP concentration increases, Mg-ADP first binds to the NBD2 consensus site to induce a conformational change of NBD2 and enhance the affinity of the NBD1 degenerate site for Mg-ADP or Mg-ATP (Fig. 6C and 6G). Then, prebound ATP together with Mg ion or newly bound Mg-ADP bridges two halves of the degenerate site to induce its full closure and NBD dimerization. Finally, there is a global conformational change of SUR1, which allosterically activates the Kir6.2 channel and occludes the GBM site of SUR1, and Kir6.2 N-terminus relocates accordingly (Fig. 6D and 6H). Mg-ATP binding at the NBD2 consensus site can have similar functions as Mg-ADP, albeit with lower affinity (Vedovato et al., 2015). Our model suggests that the essential elements for asymmetric NBD dimerization are the degenerate site of both NBDs and the consensus site of NBD2. Indeed, previous mutation results support this model. Mutations of K719A and D854N on the degenerate site of NBD1; G1479D, G1479R, G1485D, G1485R, and Q1486H on the degenerate site of NBD2; and K1385M, D1506N, and D1506A on the consensus site of NBD2 (Gribble et al., 1997; Nichols et al., 1996; Shyng et al., 1997) can impair Mg-ADP activation, whereas mutations of G827D, G827R, and Q834H on the consensus site of NBD1 can preserve Mg-ADP activation, although with altered kinetics (Nichols et al., 1996; Shyng et al., 1997). Our model also suggests that ATP hydrolysis is not required for channel activation, which is consistent with functional studies (Choi et al., 2008; Ortiz et al., 2013).

**Figure 6.**
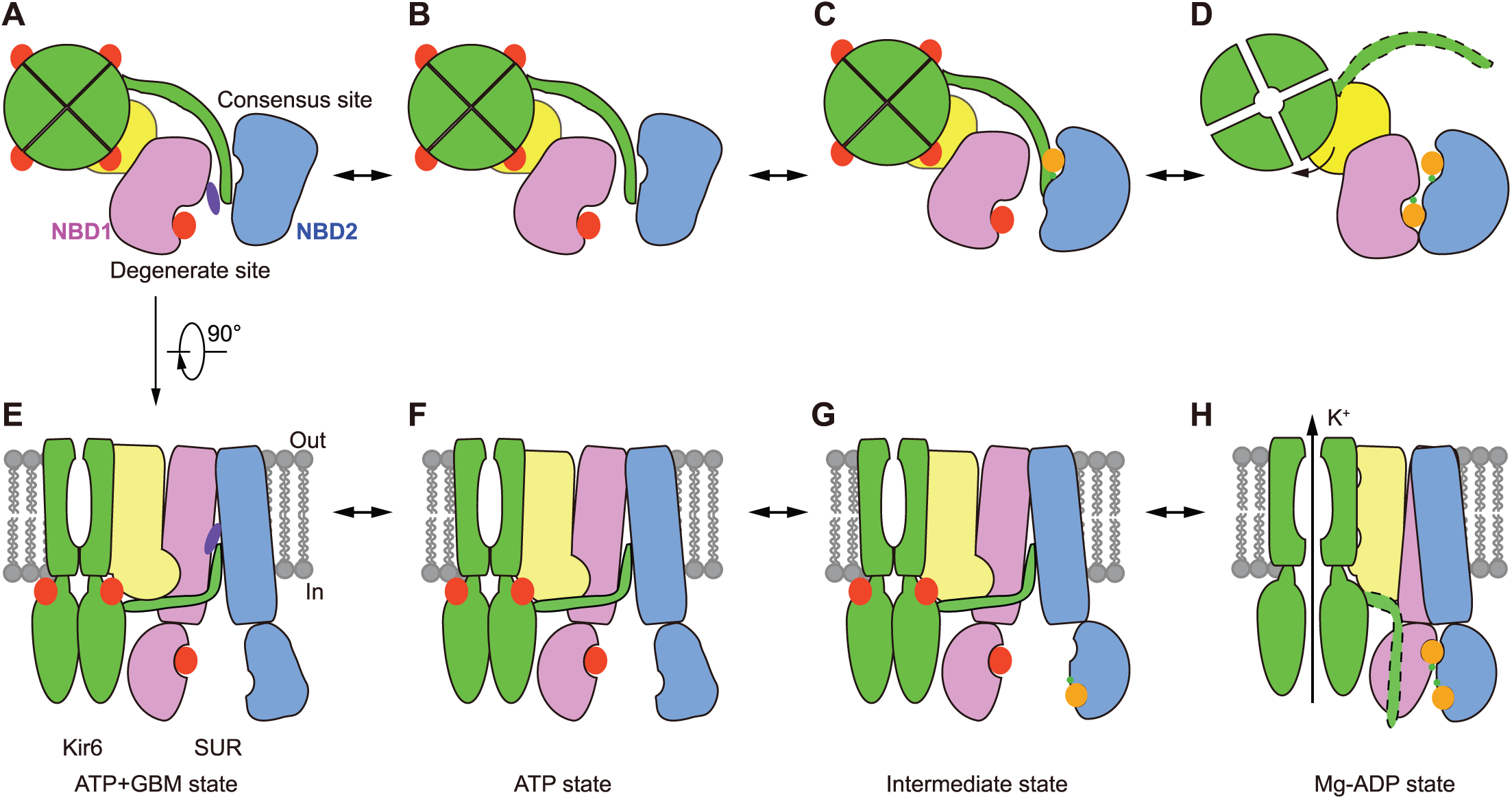
Model for nucleotide-induced NBD dimerization of the K_ATP_ channel. (A-D) Bottom view of SUR1 cartoon models from cytosolic side. For simplicity, only CTDs of Kir6.2 and TMD0-L0 and NBDs from one SUR1 subunit are shown and colored the same as in Figure 1A. KNtp with an unknown conformation in Mg-ADP state is outlined by dashed lines. ATP and ADP are shown as red and orange big spheres, respectively. Mg2+ ions are shown as green small spheres. (E-H) Side view of SUR1 cartoon models. For simplicity, a pair of Kir6.2 and one SUR1 subunit are shown.

## MATERIALS AND METHODS

### Cell Culture

Sf9 cells (Thermo Fisher Scientific) were cultured in Sf-900 III SFM medium (Thermo Fisher Scientific) at 27 °C. FreeStyle 293-F (Thermo Fisher Scientific) suspension cells were cultured in Freestyle 293 medium (Thermo Fisher Scientific) supplemented with 1% FBS at 37 °C with 6% CO_2_ and 70% humidity.

### Screening of Fusion Constructs

Fusion constructs, SUR1-Kir6.2, were created by linking the C-terminus of SUR1 from *Mesocricetus auratus* (Aguilar-Bryan et al., 1995) to the N-terminus of Kir6.2 (Woo et al., 2013) from *Mus musculus* using flexible linkers with variable lengths. The fusion constructs were cloned into a modified C terminal GFP-tagged BacMam expression vector which contains two Strep tags after GFP as described previously (Li et al., 2017). SUR1-Kir6.2 fusion constructs were screened by transient transfection of FreeStyle 293-F cells (Thermo Fisher Scientific) cultured in SMM-293TI medium (Sino Biological Inc.) using polyethylenimine (Polyscience). Cells were harvested 60 hours post-transfection and solubilized in 20 mM HEPES pH8.0 + 120 mM KCl + 1% digitonin (Biosynth) at 25°C. Cell lysates were cleared by centrifuge at 40,000 rpm for 30 min and supernatants were loaded onto Superose 6 increase column (GE Healthcare) for analysis (Kawate and Gouaux, 2006). Based on expression level and SUR1-Kir6.2 tetramer assembly, SUR1-Kir6.2 fusion with a 39-residue linker (VDGSGSGSGSAAGSGSGSGSGSGAAGSGSGSGSGSGAAA) was selected and expressed in BacMam system for further structural characterization (Goehring et al., 2014).

### Electrophysiology

K_ATP_ constructs were transfected into FreeStyle 293-F cells at density of 1×10^6^ cells/ml using polyethylenimine and cells were cultured 36-48 hours before recording. Patch electrodes were pulled from a horizontal microelectrode puller (P-1000, Sutter Instrument Co, USA) to a tip resistance of 3.0-5.0 MΩ. The pipette solution contained (mM):140 KCl, 1.2 MgCl_2_, 2.6 CaCl_2_, 10 HEPES (pH 7.4, NaOH) and the bath solution contained (mM): 140 KCl, 10 EGTA, 1 MgCl_2_, 10 HEPES (pH 7.4, NaOH). Macroscopic currents were recorded using inside-out mode at +60 mV through an Axopatch 200B amplifier (Axon Instruments, USA). Recording were started when currents were stable after patch excision. Signals were acquired at 20 kHz and lowpass filtered at 5 kHz. Data was further analyzed with pclampfit 9.0 software and filtered at 1kHz. For the GBM inhibition assay, steady state currents after GBM treatment were normalized to the currents before GBM treatment. Run-down was not corrected during data analysis.

### Protein Expression and Purification

K_ATP_ channels were expressed in FreeStyle 293-F cells as described previously (Li et al., 2017). For purification, membrane pellets were homogenized in 20 mM HEPES pH8.0 at 25°C + 120 mM KCl and then solubilized in 1% digitonin for 30 min at 4°C. Unsolubilized materials were removed by centrifugation at 40,000 rpm for 50 min in Ti45 rotor (Beckman). Supernatant was loaded onto 5 mL strep-tactin superflow high capacity resin (IBA). Resin was washed by 20 mM HEPES pH8.0 at 25°C + 120 mM KCl + 0.1% digitonin and protein was eluted by 20 mM HEPES pH8.0 at 25°C + 120 mM KCl + 0.1% digitonin + 1 mM EDTA + 5 mM desthiobiotin. Protein eluted from strep-tactin column was concentrated by 100-kDa cut-off concentrator (Millipore) and loaded onto Superose 6 increase (GE Healthcare) running in 20 mM HEPES pH8.0 at 25°C + 120 mM KCl + 0.1% digitonin. Peak fractions that contain SUR1-Kir6.2 fusion protein were combined and concentrated to A_280_ = 8 (estimated protein concentration was 9 µM K_ATP_ proteins). For cryo-EM sample preparation, the proteins were supplemented with 2 mM EDTA, 200 μM glibenclamide (TCI), and 2 mM ATPγS (Sigma) (ATP+GBM state), or 2 mM EDTA and 2 mM ATPγS (Sigma) (ATP state), or 2.5 mM MgCl_2_, 1 mM ADP (Sigma), 400 μM 08:0 PI(4,5)P_2_ (Avanti), 500 μM NN414 (Sigma), and 1 mM Na_3_VO_4_ (Mg-ADP state). Cryo-EM sample preparation was prepared as described previously (Li et al., 2017).

### Cryo-EM Data Acquisition

Cryo-grids were screened on a Talos (Thermo Fisher Scientific) operated at a voltage of 200 kV with an eagle 4k x 4k CCD camera (Thermo Fisher Scientific). Good grids were transferred into a Titan Krios (Thermo Fisher Scientific) operated at 300 kV and images were collected using K2 camera (Gatan) mounted post an quantum energy filter with 20 eV slit. K2 camera was operated under super resolution mode with super resolution pixel size 0.5275 Å (ATP+GBM state and Mg-ADP state) or 0.67 Å (ATP state) at object plane. The defocus value ranged from −1.0 μm to −3.5 μm. Data acquisitions were performed automatically using Leginon software (Suloway et al., 2005) in the movie mode with a dose rate of 3.8 or 4.7 e^-^/s/Å^2^. The total exposure was 50 e^-^/Å^2^ and each 13 or 11 seconds movie was dose-fractioned into 50 frames.

### Image Processing

Collected movies were gain-corrected, motion-corrected, exposure-filtered and binned with MotionCor2 (Zheng et al., 2017), producing dose-weighted and summed micrographs with pixel size 1.055 Å/pixel (ATP+GBM state and Mg-ADP state) or 1.34 Å/pixel (ATP state). CTF models of dose-weighted micrographs were determined using Gctf (Zhang, 2016). Autopick was done with Gautomatch (developed by Kai Zhang, MRC-LMB) using the projections of our previous map (EMD-6689) as template. Auto-picked particles were extracted from dose-weighted micrographs by binning factor of 2 and subjected to 2D classification using cryoSPARC (Punjani et al., 2017). 3D classification was done using our former K_ATP_ map (EMD-6689) as the initial model with GPU-accelerated Relion 2.0 (Kimanius et al., 2016). Particles from selected 3D classes were re-centered and re-extracted from summed micrographs without binning (1.055 Å/pixel or 1.34 Å/pixel). Re-extracted particles were subjected to 3D auto-refinement with C4 symmetry. Additional focused classification of Kir6.2 CTD with a soft mask but without alignment was used to separate 3D classes with different CTD conformations. To improve the local map quality of SUR1 ABC transporter module (maSUR1 214-1579), alignment parameters of each SUR1 ABC transporter module particles were obtained from global refined particles by symmetry expansion using relion_particle_symmetry_expand --sym C4 (Zhou et al., 2015). Then the signals outside SUR1 ABC transporter module were subtracted from each particles using particle subtraction function in Relion 2.0 as described previously (Bai et al., 2015). The signal subtracted particles were used for further 3D refinement using local search by restricting sigma_ang to 5°, using the map of SUR1 ABC transporter module as the starting model. All of the resolution estimations were based on gold standard FSC 0.143 after correction for mask effects (Chen et al., 2013). The final map was sharpened with B factor automatically determined by Relion 2.0 (Kimanius et al., 2016).

### Model Building

Individual domain of previous K_ATP_ models (PDB: 5WUA and 5TWV) and CFTR in Mg-ATP bound state (PDB: 5W81) were used as the initial model, docked into cryo-EM map with best local quality of that region by Chimera and manually rebuilt in Coot according to the density (Emsley et al., 2010). The model was further refined by phenix (Adams et al., 2010). Specifically, models of Kir6.2 transmembrane domain and SUR1 TMD0 were refined against the focused refined map of transmembrane domain in the ATP+GBM state. Models of SUR1 ABC transporter module were refined against the focused refined map in respective states. Models of Kir6.2 CTD were refined against maps after focused classification and refinement of Kir6.2 CTD in the ATP+GBM state. The individually refined models were then docked into the maps of the full K_ATP_ channel and refined respectively. Figures were prepared with Pymol (Schrödinger, LLC.) or Chimera (Pettersen et al., 2004).

### Quantification and statistical analysis

Resolution estimations of cryo-EM density maps are based on the 0.143 Fourier Shell Correlation (FSC) criterion (Chen et al., 2013).

## Author contributions

L.C. initiated the project. J.-X. W., D.D., M.W., Y.K., X.Z. and L.C. designed experiments. J.-X W. prepared the cryo-EM sample. J.-X.W., M.W., D.D., Y.K. and L.C. collected EM data. J.-X.W. and L.C. performed image processing and analyzed EM data. D.D. performed electrophysiology experiments. X.Z. performed the HPLC and mass spectrum analysis. L.C. and J.-X. W. built the model, prepared the figures and wrote the manuscript draft. All authors contributed to manuscript preparation.

## ACCESSION CODES

The 3D cryo-EM density maps have been deposited in the Electron Microscopy Data Bank under the accession number EMD: EMD-6831, EMD-6832, and EMD-6833 for the ATP+GBM state of K_ATP_, EMD-6848, EMD-6849, and EMD-6850 for the ATP state of K_ATP_, and EMD-6852, EMD-6851, and EMD-6853 for the Mg-ADP state of K_ATP_. Coordinates of the K_ATP_ structure have been deposited in the Protein Data Bank under the accession number PDB: 5YKE, 5YKF, 5YKG, 5YW7 for the ATP+GBM state of K_ATP_, 5YW8, 5YW9, and5YWA for the ATP state of K_ATP_, and 5YWC, 5YWB, and 5YWD for the Mg-ADP state of K_ATP_.

## ACKNOWLEDGMENTS

We thank all of Chen Lab members for kindly help. We also thank Joseph Bryan for sharing maSUR1 cDNA and J. Marc Simard for sharing mmKir6.2 cDNA. Cryo-EM data collection was supported by Electron microscopy laboratory and Cryo-EM platform of Peking University with the assistance of Xuemei Li and Daqi Yu, the National Center for Protein Science (Shanghai) with assistance of Liangliang Kong and Zhenglin Fu, Center for Biological Imaging, Institute of Biophysics, Chinese Academy of Science with assistance of Zhenxi Guo. Part of structural computation was also performed on the Computing Platform of the Center for Life Science and High-performance Computing Platform of Peking University.

The work is supported by grants from the Ministry of Science and Technology of China (National Key R&D Program of China, 2016YFA0502004 to L.C.) and National Natural Science Foundation of China (31622021 and 31521062 to L.C.) and Young Thousand Talents Program of China to L.C. and the China Postdoctoral Science Foundation (2016M600856 and 2017T100014 to J.-X.W.). J.-X. W. is supported by the postdoctoral foundation of the Peking-Tsinghua Center for Life Sciences, Peking University.

## ABBREVIATIONS

K_ATP_: ATP-sensitive potassium channel
cryo-EM: cryo-electron microscopy
SUR: sulfonylurea receptor
Kir6: inward-rectifying potassium channel 6
GBM: glibenclamide
ABC: ATP-binding cassettes
TMD0-L0: transmembrane domain 0-loop 0
NBD: nucleotide binding domain

## COMPLIANCE WITH ETHICS GUIDELINES

Jing-Xiang Wu, Dian Ding, Mengmeng Wang, Yunlu Kang, Xin Zeng, and Lei Chen declare that they have no conflict of interest. This article does not contain any studies with human or animal subjects performed by the any of the authors.

---

**Figure Supplement 1.**
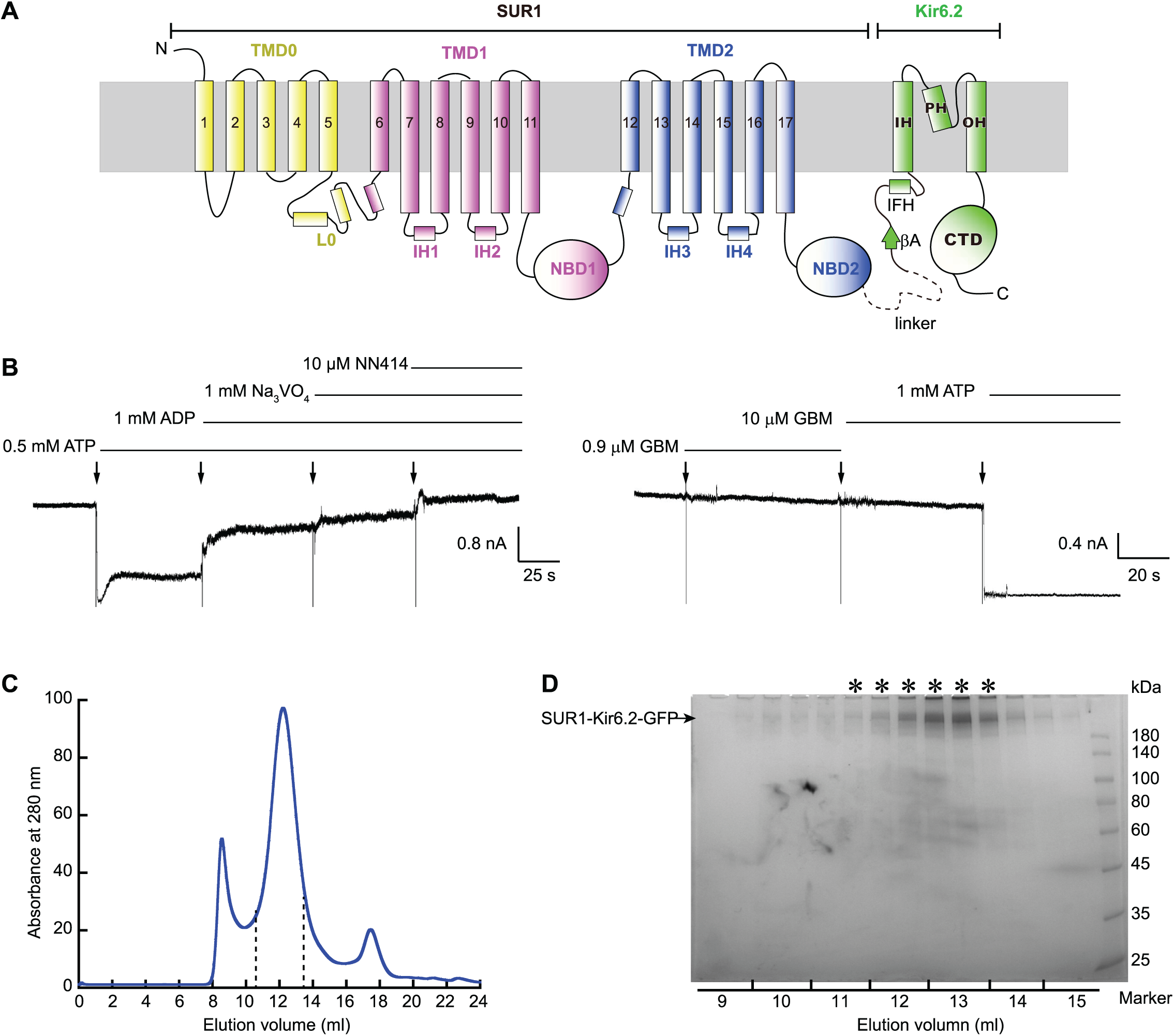
Characterization of the K_ATP_ channel complex. (A) Cartoon topology of SUR1-Kir6.2 fusion construct covalently linked by a flexible 39-residue linker (a dashed line). TMD, transmembrane domain; L0, loop 0; NBD, nucleotide-binding domain; OH, outer helix; PH, pore helix; IH, inner helix; βA, the first β strand of Kir6.2; IFH, interfacial helix; and CTD, cytoplasmic domain. (B) Activation effect of Mg-ADP and NN414 (left panel), and inhibitory effect of GBM and ATP (right panel) on the macroscopic currents of SUR1-Kir6.2 fusion construct in an inside-out mode. (C) Size exclusion chromatography (SEC) elution profiles of SUR1-Kir6.2 K_ATP_ fusion protein. Pooled fractions between dashed lines were used for cryo-EM sample preparation. (D) SDS-PAGE of purified K_ATP_ fusion protein corresponding to the indicated SEC fractions. Fractions labeled with stars were pooled and concentrated for cryo-EM grids preparation.

**Figure Supplement 2.**
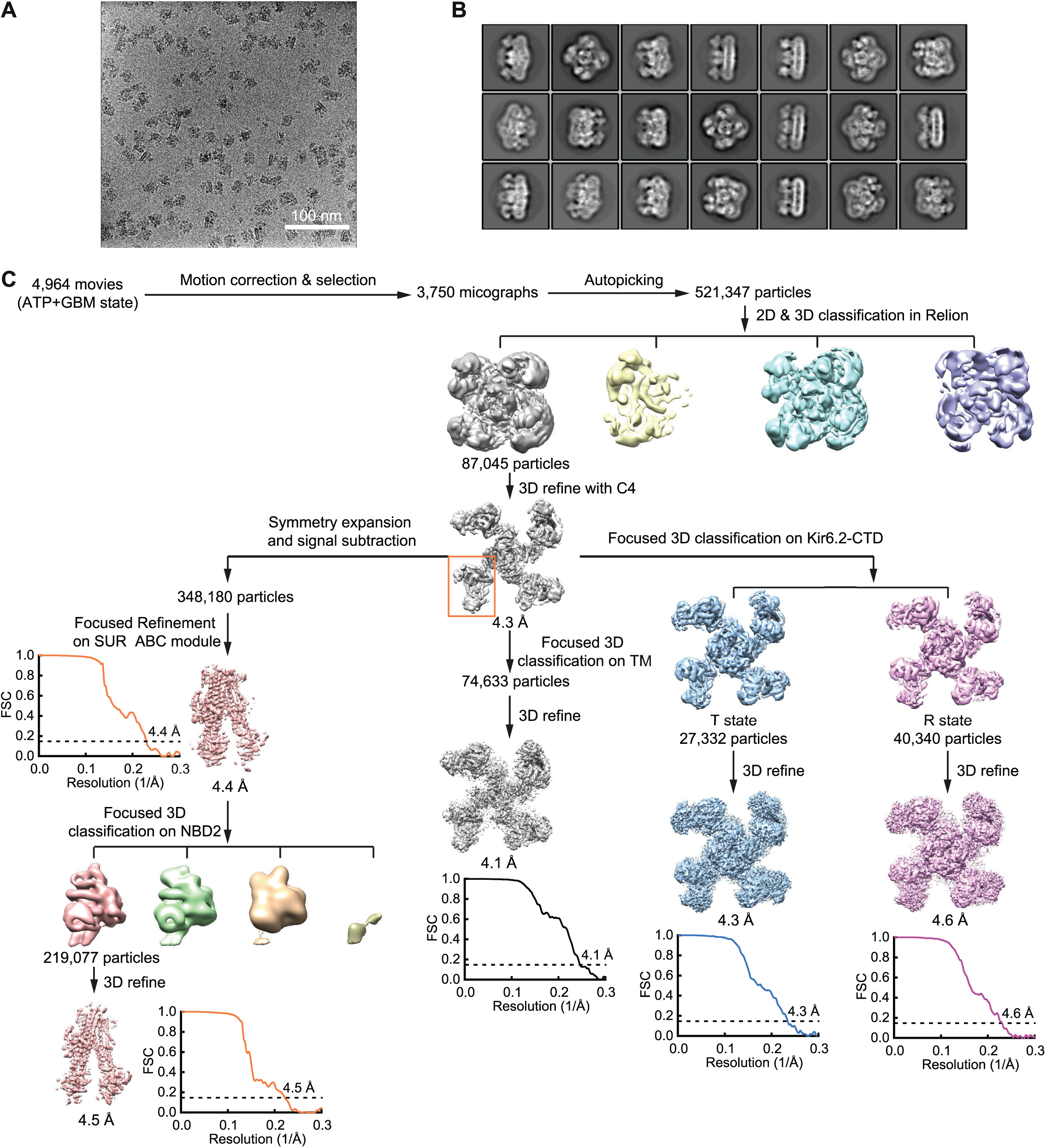
Workflow for cryo-EM data processing of K_ATP_ in complex with ATPγS and GBM (ATP+GBM state). (A) Representative raw micrograph. (B) Representative two-dimensional class averages of the K_ATP_ channel. (C) Flow-chart of EM data processing. Focused classification on Kir6.2 CTD and SUR1 NBD2, and focused refinement on SUR1 ABC transporter module were applied to improve the map quality of corresponding regions. Masked and corrected gold-standard Fourier shell correlation (FSC) curves of the final refinement are shown below the maps after post-processing. Resolution estimation was based on the criterion of FSC 0.143 cutoff.

**Figure Supplement 3.**
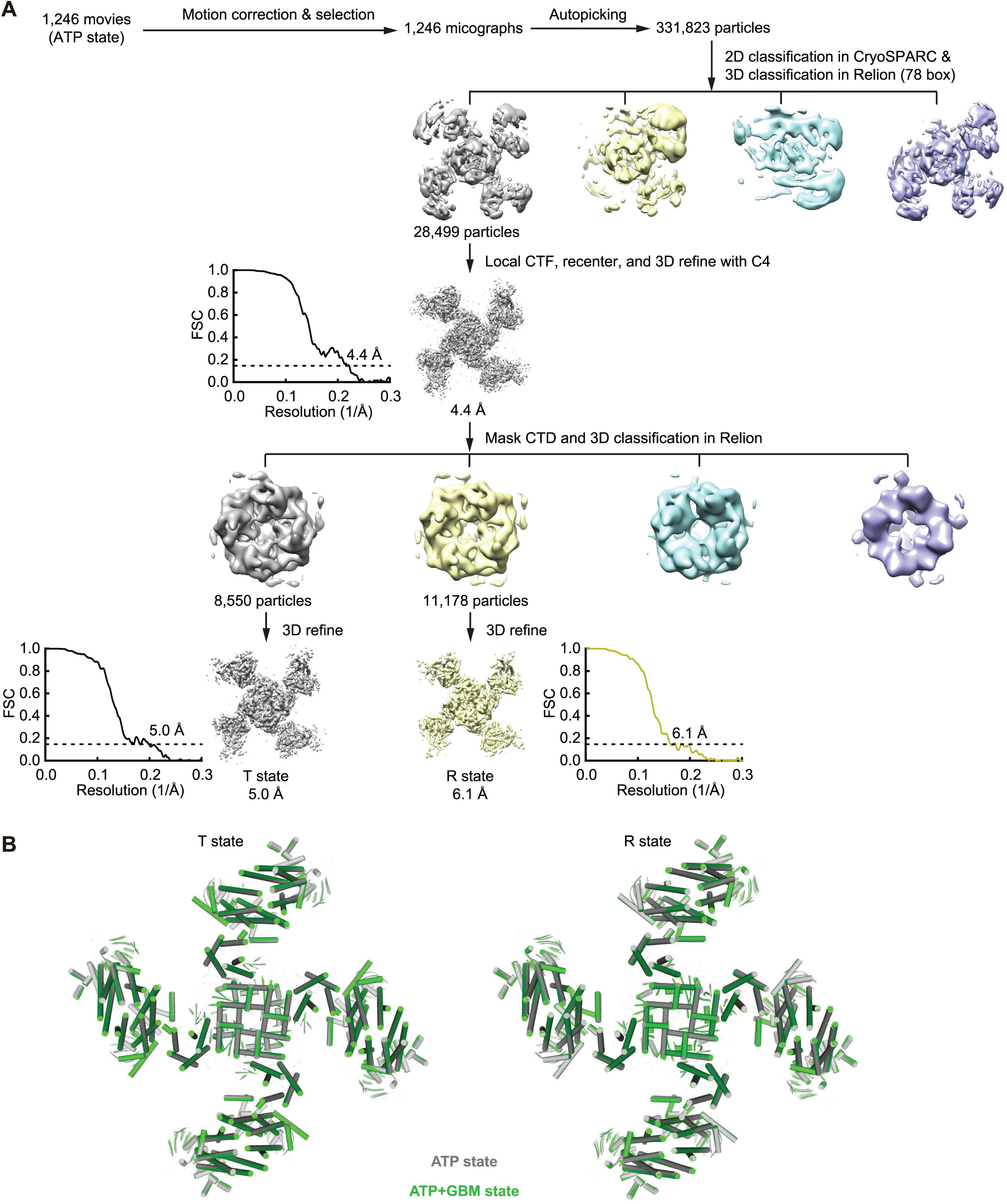
Structure of K_ATP_ in complex with ATPγS (ATP state). (A) Workflow for cryo-EM data processing of K_ATP_ in ATP state. Flow-chart of 3D classification, refinement, and focused 3D classification on Kir6.2 CTD and subsequent refinement for “T state” and “R state”. Masked and corrected gold-standard Fourier shell correlation (FSC) curves of the final refinement are shown below the maps after post-processing. Resolution estimation was based on the criterion of FSC 0.143 cutoff. (B) Top view of structural comparison of “T state” (left) and “R state” (right) structures between the ATP (green) and ATP+GBM (gray) states.

**Figure Supplement 4.**
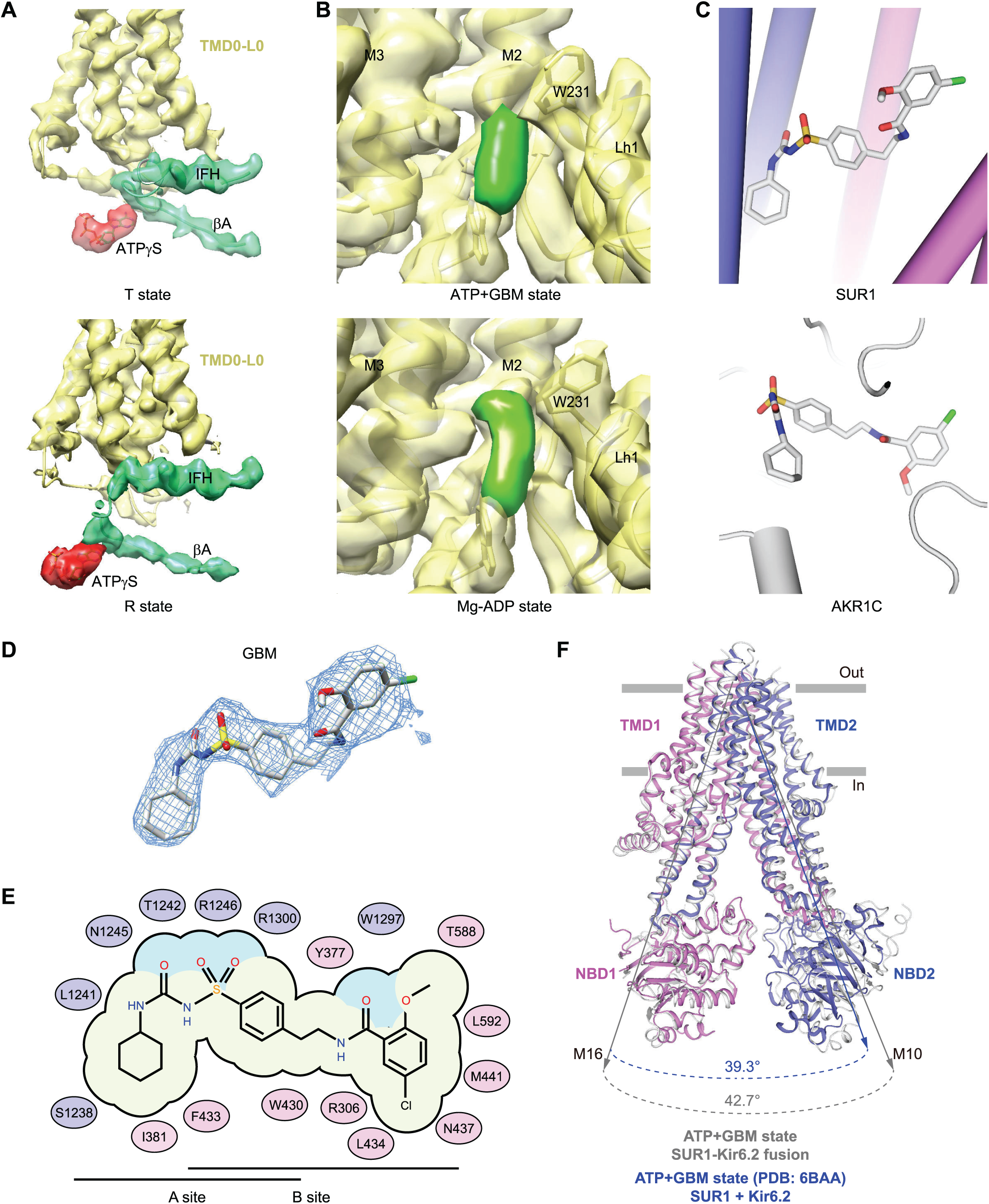
Cryo-EM map of K_ATP_ in the ATP+GBM state and the GBM binding site. (A) Local EM densities of ATPγS in “T state” (top) and “R state” (bottom) structures of the ATP+GBM state. Maps are colored the same as in Figure 1A. (B) Cryo-EM map around the lasso motif in the ATP+GBM (top) and Mg-ADP states (bottom, described later). Extra density with unknown identity is colored in green. (C) Different conformation of glibenclamide bound in K_ATP_ (top) and human AKR1C (bottom) (PDB: 4YVP). (D) Cryo-EM density of GBM. (E) Putative glibenclamide interactions with SUR1. Blue ovals represent residues on TMD1 and magenta ovals represent residues on TMD2. (F) Superposition of SUR1 ABC transporter module structures in the ATP+GBM state solved using different constructs. Structure of SUR1-Kir6.2 fusion construct in this study is shown in grey. Structure of SUR1+Kir6.2 as separated peptides (PDB: 6BAA) is colored. Transmembrane helices from the first half of TMD of SUR1 ABC transporter module are aligned. Angles between helices M10 and M16 in the two structures are shown below.

**Figure Supplement 5.**
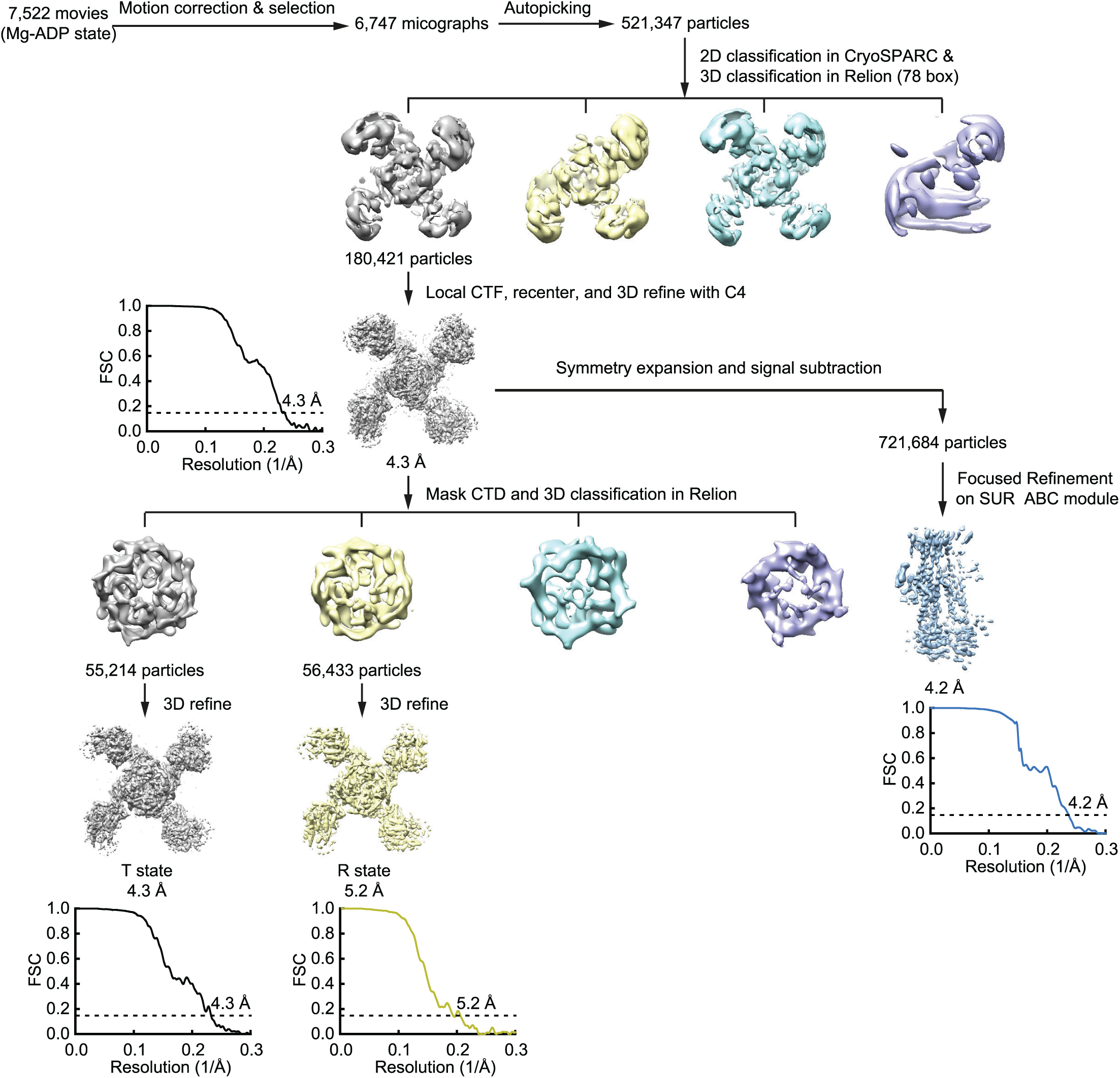
Workflow for cryo-EM data processing of K_ATP_ in the Mg-ADP state. Flow-chart of 3D classification and subsequent refinement. To improve the map quality of certain regions, focused classification and refinement were employed similarly as in Extended Data Fig. 2c. Masked and corrected gold-standard Fourier shell correlation (FSC) curves of the final refinement are shown below the maps after post-processing. Resolution estimation was based on the criterion of FSC 0.143 cutoff.

**Figure Supplement 6.**
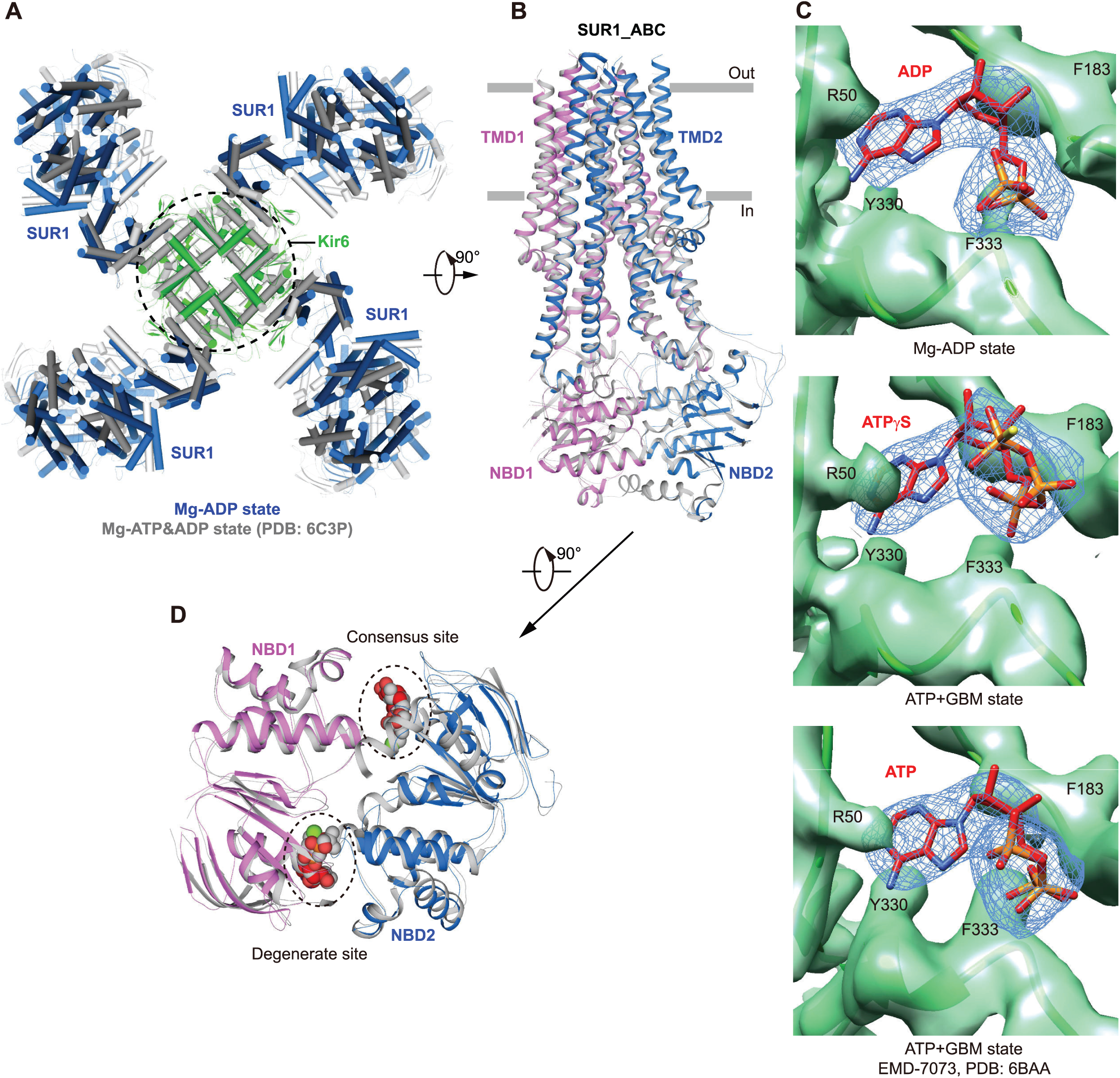
Structural comparison of K_ATP_ in different states. (A) Top view of structural comparison between the Mg-ADP (“T state”) and Mg-ATP&ADP (“propeller form”, PDB: 6C3P) states by aligning Kir6.2. (B) Superposition of the ABC module between the Mg-ADP (colored) and Mg-ATP&ADP (grey) states. (C) Local EM density maps of inhibitory site bound with ADP (top), ATPγS (middle), and ATP (EMD-7073 and PDB: 6BAA) (bottom) in different states. Maps of Kir6.2 are colored in green. Nucleotide densities are show as blue meshes with nucleotide model shown as sticks. (D) Similar asymmetric NBD dimer induced by the Mg-ADP (colored spheres) or Mg-ATP/Mg-ADP (gray spheres) molecules.

**Figure Supplement 7.**
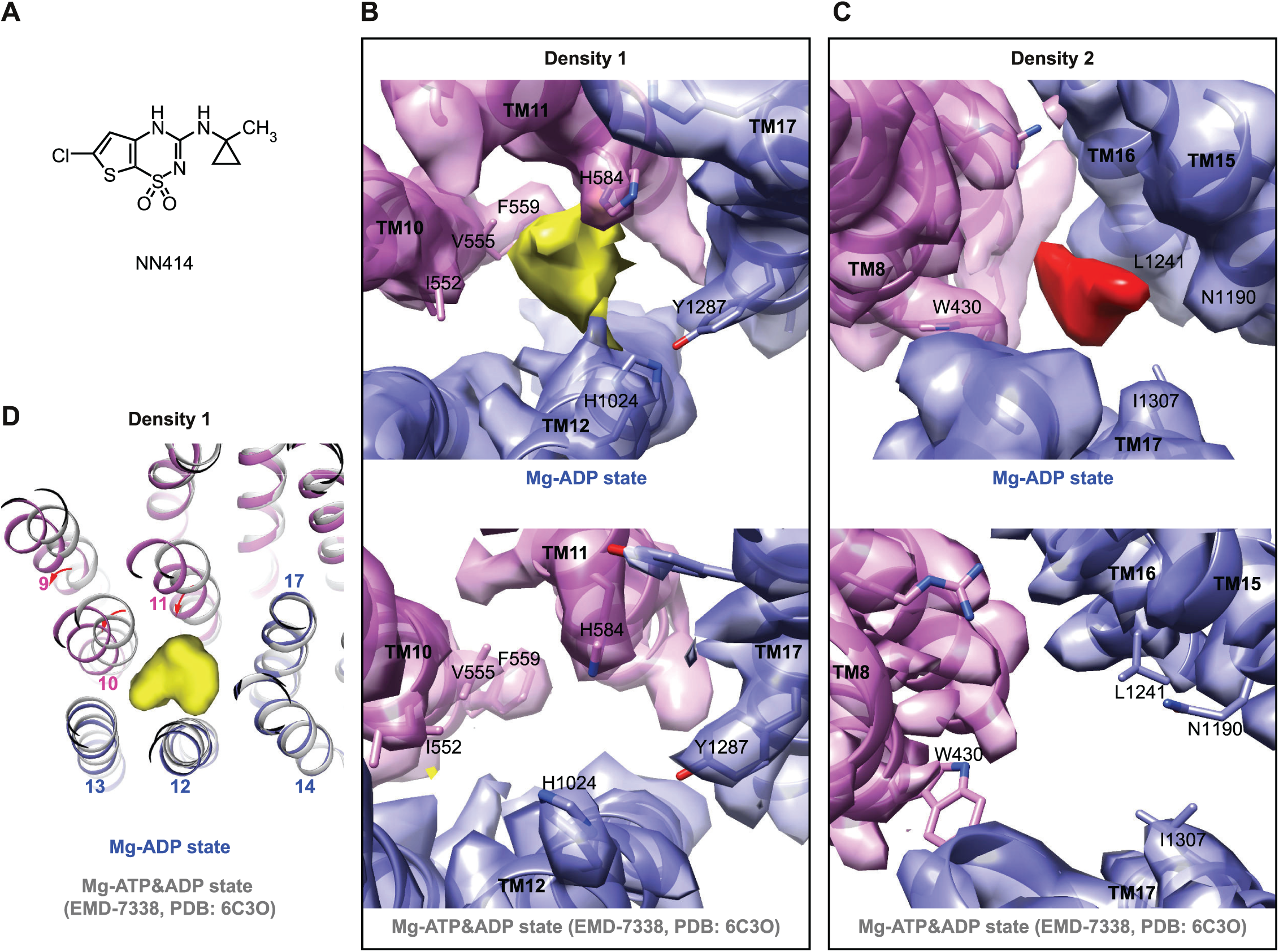
Extra densities on the map of SUR1 in the Mg-ADP state. (A) Chemical structure of NN414 (6-chloro-3-[[1-methylcyclopropyl]amino]-4H-thieno[3,2-e]-1,2,4-thiadiazine 1,1-dioxide tifenazoxide). (B) Local EM density maps surrounded by M10-12 and M17 in the Mg-ADP state (top) with one extra density (density 1: yellow). The same site in the Mg-ATP&ADP state (bottom, EMD-7338, PDB: 6C3O) is apo. (C) Local EM density maps surrounded by M8 and M15–17 in the Mg-ADP state (top) with the other extra density (density 2: red). The same site in the Mg-ATP&ADP state (bottom, EMD-7338, PDB: 6C3O) is apo. Figures are created from 4.2 Å, focused refinement map of SUR1 ABC transporter module. The cryo-EM density maps are contoured to the same level as in panel (B). Densities of SUR1 molecule are colored the same as in Figure 1. (D) Superposition of the structures with (Mg-ADP state) and without NN414 (Mg-ATP&ADP state) by aligning the TMD domain of SUR1. Local conformational changes of transmembrane helices (M9-11) at density 1 are indicated by red arrows. A surface representation of density 1 is shown in yellow.

**Table S1.** Cryo-EM data collection, refinement and validation statistics of the ATP+GBM state of K_ATP_.

|  | Focus on<br>TM | Class 1 of<br>Kir6.2 CTD | Class 2 of<br>Kir6.2 CTD | Focus on<br>SUR1 ABC<br>module | Class 1 of<br>SUR1<br>NBD2 |
| --- | --- | --- | --- | --- | --- |
|  | EMD-6831<br>5YKE | EMD-6832<br>5YKF | EMD-6833<br>5YKG | EMD-6847<br>5YW7 |  |
| **Data collection and processing** |  |  |  |  |  |
| Magnification |  |  | 47,400× |  |  |
| Voltage (kV) |  |  | 300 |  |  |
| Electron exposure (e<sup>-</sup>/Å<sup>2</sup>) |  |  | 52 |  |  |
| Defocus range (μm) |  |  | -1.5 to -3.5 |  |  |
| Pixel size (Å) |  |  | 1.055 |  |  |
| Symmetry imposed | *C4* | *C4* | *C4* | *C1* | *C1* |
| Final particle images (no.) | 74,633 | 27,332 | 40,340 | 348,180 | 219,077 |
| Map resolution (Å) | 4.1 | 4.3 | 4.6 | 4.4 | 4.5 |
| FSC threshold | 0.143 | 0.143 | 0.143 | 0.143 | 0.143 |
| Map sharpening *B* factor (Å<sup>2</sup>) | -204 | -187 | -210 | -300 | -307 |
| **Refinement** |  |  |  |  |  |
| Model composition |  |  |  |  |  |
| Non-hydrogen atoms | 30,636 | 50,988 | 50,868 | 8,944 |  |
| *B* factors (Å<sup>2</sup>) |  |  |  |  |  |
| Non-hydrogen atoms | 130.49 | 151.43 | 152.64 | 168.47 |  |
| R.m.s. deviations |  |  |  |  |  |
| Bond lengths (Å) | 0.005 | 0.005 | 0.003 | 0.003 |  |
| Bond angles (°) | 0.780 | 0.765 | 0.654 | 0.658 |  |
| Validation |  |  |  |  |  |
| MolProbity score | 2.52 | 2.71 | 2.61 | 2.57 |  |
| Clashscore | 11.31 | 15.14 | 13.19 | 13.99 |  |
| Poor rotamers (%) | 7.60 | 8.16 | 7.90 | 6.49 |  |
| Ramachandran plot |  |  |  |  |  |
| Favored (%) | 95.93 | 95.15 | 95.65 | 95.72 |  |
| Allowed (%) | 3.65 | 4.42 | 4.23 | 4.28 |  |
| Disallowed (%) | 0.42 | 0.44 | 0.12 | 0.00 |  |

**Table S2.** Cryo-EM data collection, refinement and validation statistics of the ATP state of K_ATP_.

|  | All particles<br>EMD-6848<br>5YW8 | Class 1 of Kir6.2 CTD<br>EMD-6849<br>5YW9 | Class 2 of Kir6.2 CTD<br>EMD-6850<br>5YWA |
| --- | --- | --- | --- |
| **Data collection and processing** |  |  |  |
| Magnification |  | 37,300× |  |
| Voltage (kV) |  | 300 |  |
| Electron exposure (e<sup>-</sup>/Å<sup>2</sup>) |  | 52 |  |
| Defocus range (μm) |  | -1.5 to -3.5 |  |
| Pixel size (Å) |  | 1.34 |  |
| Symmetry imposed |  | *C4* |  |
| Final particle images (no.) | 28,499 | 8,550 | 11,178 |
| Map resolution (Å) | 4.4 | 5.0 | 6.1 |
| FSC threshold | 0.143 | 0.143 | 0.143 |
| Map sharpening *B* factor (Å<sup>2</sup>) | -198 | -161 | -195 |
| **Refinement** |  |  |  |
| Model composition |  |  |  |
| Non-hydrogen atoms | 50,524 | 50,524 | 50,496 |
| *B* factors (Å<sup>2</sup>) |  |  |  |
| Non-hydrogen atoms | 219.24 | 219.24 | 505.25 |
| R.m.s. deviations |  |  |  |
| Bond lengths (Å) | 0.005 | 0.005 | 0.004 |
| Bond angles (°) | 0.796 | 0.796 | 0.693 |
| Validation |  |  |  |
| MolProbity score | 2.81 | 2.81 | 2.72 |
| Clashscore | 19.01 | 19.00 | 17.20 |
| Poor rotamers (%) | 8.11 | 8.11 | 7.91 |
| Ramachandran plot |  |  |  |
| Favored (%) | 95.13 | 95.13 | 95.65 |
| Allowed (%) | 4.43 | 4.43 | 4.17 |
| Disallowed (%) | 0.44 | 0.44 | 0.19 |

**Table S3.** Cryo-EM data collection, refinement and validation statistics of the Mg-ADP state of K_ATP_.

|  | All particles | Class 1 of<br>Kir6.2 CTD<br>EMD-6852<br>5YWC | Class 2 of<br>Kir6.2 CTD<br>EMD-6851<br>5YWB | Focus on SUR1<br>ABC module<br>EMD-6853<br>5YWD |
| --- | --- | --- | --- | --- |
| **Data collection and processing** |  |  |  |  |
| Magnification |  | 47,400× |  |  |
| Voltage (kV) |  | 300 |  |  |
| Electron exposure (e<sup>-</sup>/Å<sup>2</sup>) |  | 52 |  |  |
| Defocus range (μm) |  | -1.5 to -3.5 |  |  |
| Pixel size (Å) |  | 1.055 |  |  |
| Symmetry imposed | *C4* | *C4* | *C4* | *C1* |
| Final particle images (no.) | 180,421 | 55,214 | 56,433 | 721,684 |
| Map resolution (Å) | 4.3 | 4.3 | 5.2 | 4.2 |
| FSC threshold | 0.143 | 0.143 | 0.143 | 0.143 |
| Map sharpening *B* factor (Å<sup>2</sup>) | -227 | -188 | -265 | -269 |
| **Refinement** |  |  |  |  |
| Model composition |  |  |  |  |
| Non-hydrogen atoms |  | 51,108 | 50,928 | 8,997 |
| *B* factors (Å<sup>2</sup>) |  |  |  |  |
| Non-hydrogen atoms |  | 283.99 | 300.02 | 203.33 |
| R.m.s. deviations |  |  |  |  |
| Bond lengths (Å) |  | 0.004 | 0.004 | 0.003 |
| Bond angles (°) |  | 0.796 | 0.776 | 0.718 |
| Validation |  |  |  |  |
| MolProbity score |  | 2.69 | 2.62 | 2.49 |
| Clashscore |  | 14.70 | 13.27 | 12.19 |
| Poor rotamers (%) |  | 7.97 | 7.66 | 6.20 |
| Ramachandran plot |  |  |  |  |
| Favored (%) |  | 95.26 | 95.50 | 95.87 |
| Allowed (%) |  | 4.47 | 4.07 | 4.13 |
| Disallowed (%) |  | 0.37 | 0.43 | 0.00 |

